# ATRX binds to atypical chromatin domains at the 3’ exons of ZNF genes to preserve H3K9me3 enrichment

**DOI:** 10.1101/027789

**Authors:** David Valle-García, Zulekha A. Qadeer, Domhnall S. McHugh, Flávia G. Ghiraldini, Asif H. Chowdhury, Dan Hasson, Michael A. Dyer, Félix Recillas-Targa, Emily Bernstein

## Abstract

ATRX is a SWI/SNF chromatin remodeler proposed to govern genomic stability through the regulation of repetitive sequences such as rDNA, retrotransposons, and pericentromeric and telomeric repeats. However, few direct ATRX target genes have been identified and high-throughput genomic approaches are currently lacking for ATRX. Here we present a comprehensive ChIP-sequencing study of ATRX in multiple human cell lines, in which we identify the 3’ exons of zinc finger genes (ZNFs) as a new class of ATRX targets. These 3’ exonic regions encode the zinc finger motifs, which can range from 1-40 copies per ZNF gene and share large stretches of sequence similarity. These regions often contain an atypical chromatin signature: they are transcriptionally active, contain high levels of H3K36me3 and are paradoxically enriched in H3K9me3. We find that these ZNF 3’ exons are co-occupied by SETDB1, TRIM28 and ZNF274, which form a complex with ATRX. CRISPR/Cas9-mediated loss-of-function studies demonstrate (i) a reduction of H3K9me3 at the ZNF 3’ exons in the absence of ATRX and ZNF274 and, (ii) H3K9me3 levels at atypical chromatin regions are particularly sensitive to ATRX loss compared to other H3K9me3-occupied regions. As a consequence of ATRX or ZNF274 depletion, cells with reduced levels of H3K9me3 show increased levels of DNA damage, suggesting that ATRX binds to the 3’ exons of ZNFs to maintain their genomic stability through preservation of H3K9me3.

## INTRODUCTION

Chromatin remodeling proteins act through shifting, sliding, deposition and eviction of nucleosomes and histones. Members of the SWI/SNF family of chromatin remodelers are fundamental in many cellular processes such as transcription, replication, DNA repair and recombination. ^1–5^ One notable chromatin remodeler involved in all of these processes is ATRX. Increasing evidence supports that ATRX acts as a sentinel of genome integrity by maintaining heterochromatin at repetitive sequences. ^6,7^ Interestingly, ATRX germline mutations are responsible for a complex genetic disorder called ATR-X (<u>A</u>lpha <u>T</u>halassemia, Mental <u>R</u>etardation <u>X</u>-linked) syndrome while somatic mutations, deletions, and altered ATRX expression levels are highly prevalent in a wide variety of cancers. ^8,9^

ATRX contains two highly conserved domains: the ADD (ATRX-DNMT3-DNMT3L) and the SWI/SNF helicase domain. ^10^ The ADD domain contains a PHD finger that binds H3K9me3/H3K4me0, ^11,12^ whereas the SWI/SNF domain is an ATP-dependent helicase responsible for the chromatin remodeling capacity of ATRX. ^10,13^ Despite the fact that ATRX binds H3K9me3/H3K4me0 *in vitro*, ATRX binds to only a subset of H3K9me3-containing regions *in vivo*. ^11,11,14,15^ In particular, ATRX is highly enriched at certain H3K9me3-containing repetitive regions such as telomeric and pericentromeric repeats as well as some retrotransposon families. ^16–22^ Furthermore, ATRX physically interacts with other H3K9me3 binding proteins such as HP1α. ^23,24^ Altogether, these pieces of evidence suggest that ATRX is involved in the regulation of particular H3K9me3-modified chromatin.

A well-characterized role of ATRX is deposition of histone variants into the chromatin template. For example, ATRX and DAXX (death-domain associated protein) act together as a histone chaperone complex for the H3 variant H3.3. ATRX is required for the localization of H3.3 at telomeres and pericentromeric repeats, ^16–19^ retrotransposons ^20–22^ and imprinted loci, ^25^ which all contain H3K9me3. This ability appears to be unique for ATRX, as the HIRA complex deposits H3.3 only at euchromatic regions. ^17,26–28^ In addition to promoting H3.3 deposition, our group showed that ATRX negatively regulates the deposition of histone variant macroH2A at the α-globin locus. ^29^

ATRX has also been implicated in resolving aberrant secondary DNA structures, called G-quadruplexes, which form in guanine-rich regions during replication and transcription. ^15,30,31^ G-quadruplexes are a common feature of some families of repetitive sequences and tandem repeats, such as those found in telomeres. Intriguingly, ATRX mutations in cancer have been linked to the Alternative Lengthening of Telomeres (ALT) pathway. ^32–35^ Although the precise role of ATRX in ALT remains unclear, it has been suggested that ATRX prevents Homologous Recombination (HR) between telomeric sequences through the resolution of stalled replication forks in G-rich regions. ^6,36^ In accordance with its role as a regulator of genome stability, several reports demonstrated that ATRX depletion causes telomere dysfunction, increased replication fork stalling and increased sensitivity to replicative stress across different cellular and *in vivo* models. ^18,22,37–40^

Despite these important functions, surprisingly few direct ATRX target genes have been identified. To address this, we utilized an unbiased approach using the ENCODE Tier 1 human erythroleukemic cell line K562 as a model system to analyze ATRX genomic occupancy. Through comprehensive ChIP-seq analyses, we identified an unexpected binding pattern of ATRX at the 3’ exons of Zinc Finger Genes (ZNFs). ZNFs represent the largest family of putative transcription factors in the human genome with more than seven hundred identified members. ^41–43^ This enrichment of ATRX at ZNF 3’ exons was further confirmed in additional human cell lines of both normal or cancer origin.

The 3’ exons of ZNFs are enriched in chromatin that is permissive to transcription yet contains high levels of H3K9me3 and H3K36me3. ^44^ These atypical chromatin regions do not possess the characteristics of any known regulatory region (i.e. promoter, enhancer, insulator) and their functional significance remains unclear. ^44^ Here we show that ATRX co-occupies 3’ ZNF exons containing an H3K9me3/H3K36me3 chromatin signature, together with the H3K9 methyltransferase SETDB1 (also known as ESET), the co-repressor TRIM28 (also known as KAP1), and the transcription factor ZNF274. Deletion of ATRX or ZNF274 leads to a reduction of H3K9me3, particularly at 3’ ZNF exons and other H3K9me3/H3K36me3-containing regions, as well as increased DNA damage, and defects in the cell cycle. Taken together, our studies suggest that ATRX binds the 3’ exons of ZNFs to maintain genomic stability by regulating H3K9me3 levels.

## RESULTS

### ATRX binds to the 3’ exons of ZNF genes <u>in K562 cells</u>

In order to perform an unbiased search for novel direct ATRX target genes, we examined its genomic distribution by ChIP-seq analysis in the human erythroleukemic cell line K562 using two independent antibodies (see Methods for details). We chose K562 as a model system for two reasons: first, it has been established that ATRX has important roles in the regulation of the erythroid lineage; ^15,29^ second, K562 is a Tier 1 ENCODE cell line that has been extensively analyzed using a wide array of genomic and epigenomic methodologies that are publicly available. ^45^

To determine the global ATRX binding pattern in relation to other chromatin modifications, we reanalyzed the available ChIP-seq ENCODE datasets for K562 (see Methods) and performed a correlation analysis of their binding profiles. The only datasets that show a positive correlation with ATRX are H3K9me3 (r = 0.46, spearman correlation) and macroH2A (r = 0.19, spearman correlation) (**Fig. S1A**), consistent with its role as an H3K9me3 binding protein ^11,12^ and macroH2A regulator, respectively. ^29,46,47^ Furthermore, we examined the genomic distribution of ATRX significant peaks and found that, consistent with previous reports, ^15,20–22^ ATRX is bound mainly to repetitive sequences (~56% of ATRX peaks overlap with repeats) (**Fig. S1B**). In order to understand ATRX distribution at a functional level, we analyzed its distribution across Hidden Markov Model-derived chromatin states. ^48^ While ATRX binds to repressed and repetitive regions (**Fig. 1A**), it is significantly enriched in transcribed regions as well (**Fig. 1A**). In order to further investigate the <u>functional</u> significance of ATRX occupancy at these transcriptionally active regions, we performed Gene Ontology (GO) analysis with

**Figure 1.**
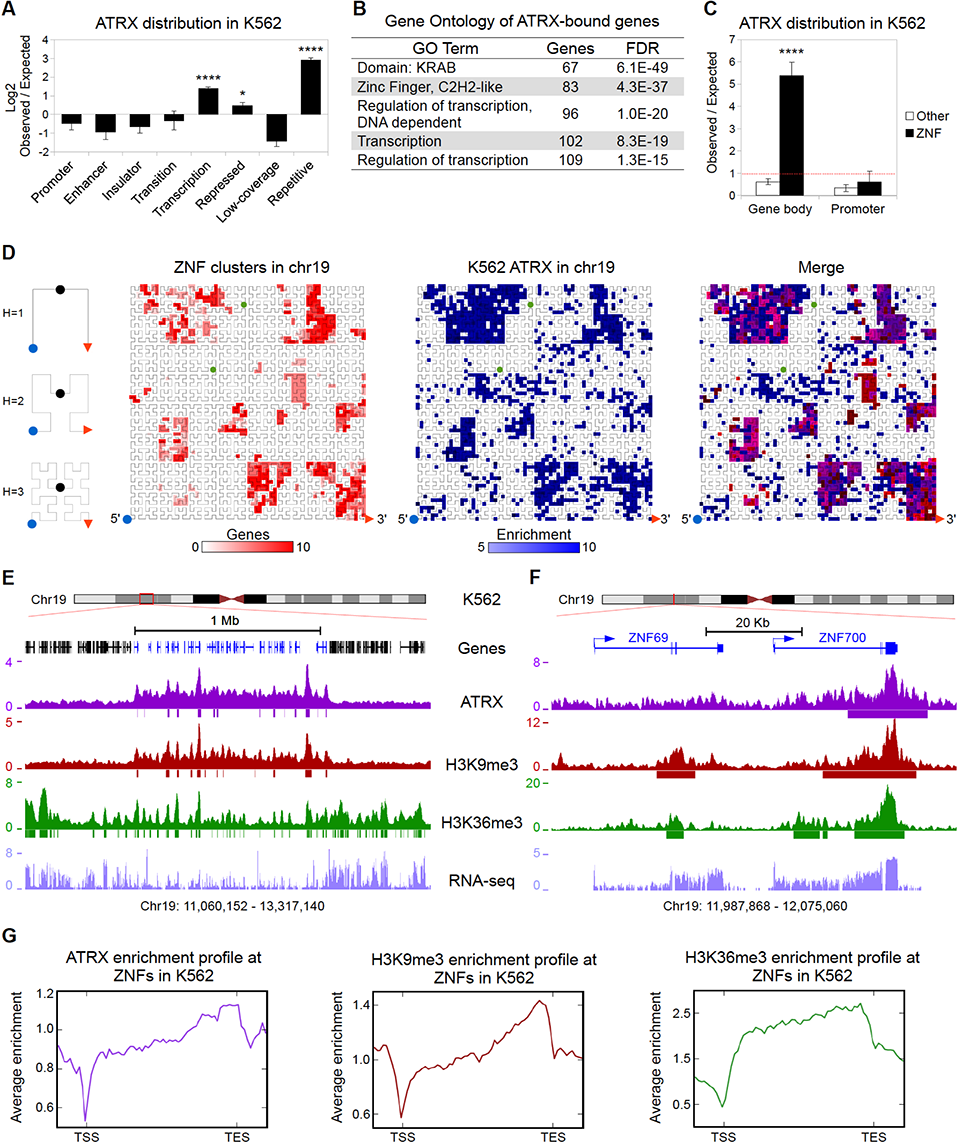
ATRX binds to the 3’ regions of ZNF genes <u>in K562 cells</u>. (A) Observed over expected random distribution of significant ATRX peaks within HMM chromatin categories <u>in K562</u>. ^48^ In (A) and (C) error bars represent standard deviation. Asterisks represent significantly overrepresented regions (* pvalue < 0.05; **** pvalue < 1×10^−4^) assessed by the hypergeometric test (see **Tables S2 and S3** for details of statistical tests). (B) Gene Ontology analysis of genes that overlap with significant ATRX peaks in their gene bodies. (C) Observed over expected random distribution of significant ATRX peaks in ZNFs and non-ZNF (other) promoters and gene bodies <u>in K562</u>. The red line represents the expected value of a random distribution. (D) Hilbert curve plot of chromosome 19 showing the ZNF clusters (red, left), highly enriched ATRX regions in <u>K562</u> (fold enrichment over input >5, blue, middle), and overlap (right). The blue dot and the red arrow mark the start (5’) and end (3’) of the chromosome, respectively (see diagram on left). The green dots indicate the start and end of centromere, which is excluded from the analysis. (E) UCSC Genome browser screenshot of a typical ZNF cluster (genes in blue) on chromosome 19. The enrichment over input signal <u>in K562</u> for ATRX, H3K9me3 and H3K36me3 ChIP-seq is shown. Significant peaks represented as bars below enrichment tracks. Normalized RPKM signal for RNA-seq is also shown. (F) Zoomed in snapshot of two ZNFs contained within the ZNF cluster shown in (E). (G) Average <u>K562</u> enrichment ChIP-seq profiles of ATRX, H3K9me3 and H3K36me3 over all ZNF gene bodies +/-1kb (n = 736).

ATRX-bound genes (n=374, **Table S1**). Strikingly, C2H2 Zinc Finger genes (ZNFs) were the most overrepresented gene family and comprised one quarter of the ATRX-bound genes, many of which contain the repressive KRAB domain (**Fig. 1B**). We next analyzed ATRX binding at promoters and gene bodies and found that enrichment of ATRX at gene bodies of ZNFs was highly significant as compared to non-ZNF genes, but that promoter regions had minimal binding in either group of genes (**Fig. 1C**).

ZNFs represent the fastest expanding gene family in the primate lineage. Frequent gene duplications and rapid divergence of paralogs are characteristic of ZNFs. ^41–43^ Because of this, ZNFs are often arranged in large continuous clusters in the human genome and share stretches of highly similar DNA sequence, particularly at their 3’ exons where the DNA sequence encoding the zinc finger motifs is contained. ^41,42^ Chromosome 19 contains the majority of ZNF clusters in the human genome. ^41–43^ By examining ATRX enrichment on chromosome 19, we found that the ZNFs clusters are demarcated by ATRX occupancy (**Fig. 1D, E**). We next analyzed the binding pattern of ATRX over individual ZNF genes and found ATRX to be preferentially enriched at the 3’ exons of ZNFs (**Fig. 1F, G**). These results were confirmed by ChIP-seq with a second ATRX antibody, which showed nearly identical enrichment patterns at ZNF genes (**2**). Overall, our ChIP-seq studies demonstrate that ZNFs are a novel set of ATRX targets and that ATRX is highly enriched at their 3’ exons.

### ATRX is enriched at ZNF genes harboring an atypical chromatin signature and distinctive epigenetic and genomic features

A large proportion of ZNFs contain an atypical chromatin signature at their 3’ exons. ^44^ This includes high levels of H3K9me3, permissibility to transcription, and enrichment of H3K36me3, a mark associated with transcriptional elongation. Our analysis of ENCODE ChIP-seq data <u>in K562 cells</u> corroborated these observations (**Fig. 1E–G**).

To investigate the epigenetic and genetic characteristics of the ATRX-enriched ZNFs and their relationship with the above atypical chromatin signature, we categorized all ZNF genes into three classes based on their ATRX enrichment levels: Class I represents ZNFs highly enriched for ATRX, Class II contains ZNFs moderately enriched, and Class III for ZNFs depleted of ATRX enrichment (**Fig. 2A**, top). We next quantified the average ChIP-seq signals of ATRX, H3K9me3 and H3K36me3 over the gene bodies of the ZNF classes. As shown in **Figure 2A**, Class I ZNFs show high levels of both H3K9me3 and H3K36me3. In contrast, Class III genes are largely depleted of H3K9me3 and show less enrichment of H3K36me3. To analyze if ATRX enrichment is correlated with the presence of H3K9me3 and H3K36me3 at the same loci, we calculated the Spearman correlation of these marks in Class I and Class III ZNFs. H3K9me3 is correlated with ATRX and moderately correlated with H3K36me3 in Class I ZNFs whereas a poor correlation for both marks was observed in Class III ZNFs (**Fig. 2B**). These data suggest that ATRX is specifically enriched at those ZNFs displaying an atypical chromatin signature.

**Figure 2.**
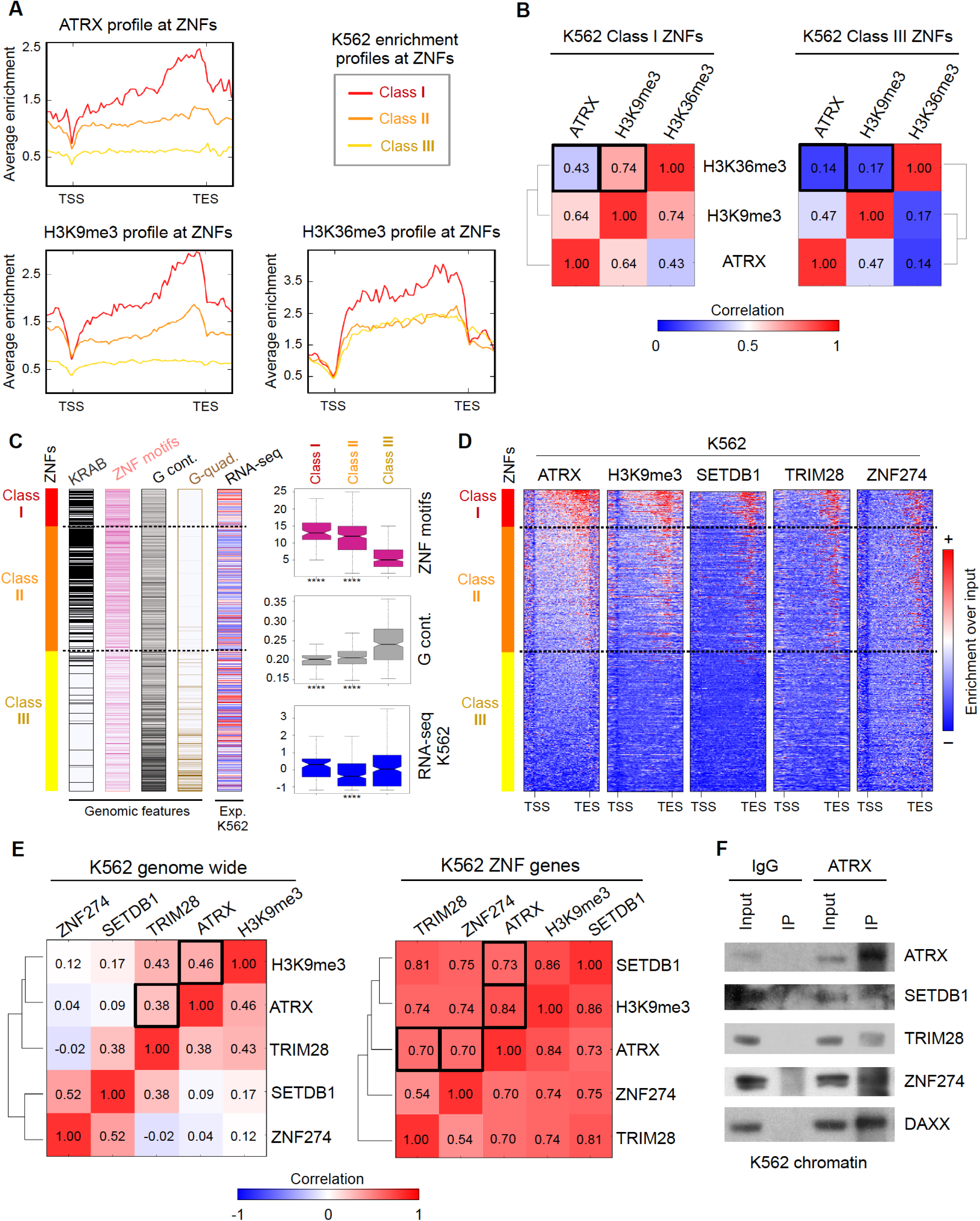
ATRX and the ZNF274/TRIM28/SETDB1 complex bind to ZNF genes with an atypical chromatin signature and distinctive genomic and epigenetic features. (A) Average <u>K562</u> enrichment ChIP-seq profiles of ATRX, H3K9me3 and H3K36me3 at ZNFs classified by their ATRX content. Class I contains high levels of ATRX (n = 91), Class II contains medium to low levels of ATRX (n = 303) and Class III is devoid of ATRX enrichment (n = 342). (B) Spearman correlation heatmap of <u>K562</u> ChIP-seq signal at ZNF Class I genes (left) and ZNF Class III genes (right). (C) Left: Distribution of genetic features among ZNF classes (sorted from high to low ATRX enrichment from top to bottom). Dashed lines show separation of the three classes. Colors represent presence of KRAB domains (black), number of zinc finger motifs (pink), G content at the C-terminal ZNF region (last 3kb of the gene) (gray) and presence of sequences predicted to form G-quadruplexes (brown). RNA-seq bar shows the Z score of the normalized RPKM signal (log2(RPKM*1)) <u>in K562</u>; red = high expression signal and blue = low expression signal. For statistical tests between the classes see **Table S3**. Right: Box plots displaying the number of ZNF motifs, G-content at the ZNF region and RNA-seq values <u>in K562</u> per ZNF Class. Asterisks show significant differences (pvalue < 1×10^−4^). (D) Metagene analysis of ChIP-seq enrichment over input profiles at ZNF gene bodies +/-1kb. (E) Spearman correlation heatmaps between ChIP-seq profiles genomewide (left) and at ZNF genes (right). Black boxes indicate the significant correlations. (F) Immunoblots for endogenous ATRX Co-IP of chromatin bound proteins in K562 cells after pulldown with IgG or ATRX antibody. DAXX used as a positive control for the ATRX IP.

To analyze the relation between ATRX and atypical chromatin genome-wide, we reanalyzed ENCODE H3K36me3 and H3K9me3 ChIP-seq data for K562 cells and plotted ATRX enrichment levels at atypical chromatin <u>peaks</u> (overlapping H3K9me3/H3K36me3 peaks), H3K9me3-only peaks and H3K36me3-only peaks. ATRX is enriched at a subset of atypical and H3K9me3-only regions, and largely absent of H3K36me3-only regions, suggesting that ATRX recruitment is independent of the H3K36me3 mark (**Fig. S3A,B**). Interestingly, ATRX enrichment is more significant at atypical chromatin regions than at

H3K9me3-only regions (**Fig. S3C**). These data suggest that ATRX is a bona fide binder of atypical chromatin genome wide.

To further understand ATRX recruitment and function at ZNFs, we analyzed the ZNF-related genomic features in the three defined classes. ZNF genes can be classified as transcriptional activators or KRAB-containing repressors. KRAB is a potent repressor domain that is contained in about half of the ZNFs and is generally encoded in two exons independent of the 3’ exon containing the zinc finger motifs. ^42^ As KRAB-containing genes were enriched in our Gene Ontology analysis of ATRX-bound genes (**Fig. 1B**), we plotted the number of KRAB domains contained by each ZNF ordered by class. Most of the ATRX-enriched Class I and Class II ZNFs contained KRAB domains, while very few of Class III ZNF genes contain this feature (**Fig. 2C**, left).

Because the DNA sequence encoding C2H2 zinc finger motifs is similar between ZNFs, it has been proposed that ZNF genes are prone to homologous recombination (HR), particularly those with more zinc finger motifs. ^41,42,44^ Therefore, the presence of H3K9me3 at the ZNF 3’ exons has been suggested to protect against HR. ^44,49^ To support this idea, we plotted the number of predicted C2H2 zinc finger motifs per ZNF gene. On average, human ZNF genes contained ~9 zinc finger motifs per gene. In contrast, Class I ZNF genes contained significantly more motifs with an average of ~14, while ATRX depleted Class III genes contained only ~6 domains per gene (**Fig. 2C**, left and top right). These results suggest that ATRX enrichment at the ZNF 3’ exons positively correlates with the number of C2H2 zinc finger motifs. This is in accordance with a study that reported H3K9me3 enrichment at 3’ ZNF exons positively associated with the number of zinc finger motifs. ^44^

A genomic feature proposed to be important for ATRX binding is the Guanine DNA content (G-content). ATRX binds to G-quadruplexes with high affinity *in vitro* and facilitates polymerase elongation through deposition of H3.3 specifically in G-rich regions that have a tendency to form these structures. ^15,31^ Based on these observations, we measured the G-content and predicted the potential of G-quadruplex formation at the 3’ ends of the ZNF genes. Surprisingly, ATRX enrichment levels negatively correlated with both G-content and the potential to form G-quadruplexes (**Fig. 2C**, left and middle right). This strongly indicates that ATRX recruitment to ZNF 3’ exons is not mediated by its ability to recognize G-quadruplexes, but by an alternative mechanism(s).

We then investigated whether ATRX enrichment and the presence of the atypical chromatin signature correlates with ZNF transcriptional levels. Thus, we analyzed the ENCODE RNA-seq datasets for K562 and plotted the normalized RPKM signal for the three ZNFs classes. Intriguingly, there was no evident association between RNA-seq expression levels and ATRX enrichment. This suggests that neither ATRX binding nor the formation of the atypical chromatin signature have a direct effect on ZNF expression levels (**Fig. 2C**, left and bottom right).

As ATRX regulates late stalled replication forks and H3K9me3-marked chromatin is often late replicating, ^37,39,40,50^ we queried whether ATRX binds to late-replicating ZNFs. To address this, we analyzed the K562 Repli-seq data from ENCODE and quantified the signal for the ZNF classes throughout S phase. Interestingly, we found that ZNF Classes I and II tend to be late replicating while Class III ZNFs replicate early (**Fig. S4A**).

In summary, we have established that ATRX levels positively correlate with H3K9me3 at atypical chromatin found at the 3’ of ZNF genes. ATRX enrichment at ZNFs is independent of transcriptional levels. Moreover, ATRX-enriched ZNFs tend to be late replicating, contain KRAB domains, and have a large number of zinc finger motifs. Such ZNFs also contain low levels of G-content and low potential for G-quadruplex formation. These trends are all statistically significant (**Tables S2-S3**).

### SETDB1, TRIM28 and ZNF274 co-iocaiize at 3’ exons of ATRX-bound ZNF genes and form a complex with ATRX

In order to find additional chromatin factors that bind the ZNF 3’ exons, we performed metagene analyses at the ZNF gene bodies with all available K562 ChIP-seq datasets from the ENCODE project. ^51^ From the 98 datasets we analyzed, we found that the enrichment levels of the H3K9me3 methyltransferase SETDB1 (also known as ESET) and the SETDB1-interacting protein TRIM28 (also known as KAP1) correlated appreciably with ATRX and H3K9me3 at the 3’ exons of ZNF genes (**Fig. 2D**). <u>Interestingly,</u> TRIM28 is a co-repressor that interacts with KRAB-containing ZNF transcription factors and recruits HDACs and H3K9 methyltransferases to enforce silencing. ^52^ Moreover, we found a striking co-localization of ATRX and H3K9me3 with the KRAB-containing transcription factor ZNF274 (**Fig. 2D**). This is in agreement with a study that reported ZNF274 to bind the 3’ region of ZNF genes and recruit SETDB1 through its interaction with TRIM28. ^53^

We then performed a correlation analysis of the ChIP-seq signals of ATRX, ZNF274, TRIM28, SETDB1 and H3K9me3 to determine if they preferentially co-localize at ZNFs. Genome-wide, ATRX moderately associates only with H3K9me3 and TRIM28 (**Fig. 2E**, left). In striking contrast, when focusing only on the ZNF genes, the correlation coefficients between ZNF274/TRIM28/SETDB1/ATRX denote a strong association, along with presence of H3K9me3 signal (**Fig. 2E**, right).

To investigate whether the ZNF274/TRIM28/SETDB1 complex interacts with ATRX, we performed co-immunoprecipitation experiments of chromatin-bound proteins in K562 cells. As show in **Figure 2F**, ATRX is able to co-immunoprecipitate with all three factors. As a control for ATRX pull-down, we also confirmed ATRX’s interaction with DAXX in K562. This data strongly suggests that ATRX physically interacts with ZNF274/TRIM28/SETDB1 at the chromatin level.

Finally, we performed a comprehensive motif analysis of ZNF274 binding sites (see Methods). We found 3 significant DNA motifs that were highly enriched at the 3’ end of ATRX enriched Class I and Class II ZNF genes, but not at ATRX depleted Class III ZNFs (**Fig. S4B**). These data favors the idea that the ZNF274/TRIM28/SETDB1/ATRX complex could bind to ZNF genes at least in part through the recognition of ZNF274binding motifs. Of note, the motifs that we found are in accordance with a previous ZNF274 motif-analysis. ^53^

Collectively, we have identified a ZNF274/TRIM28/SETDB1/ATRX complex that localizes to the 3’ region of ZNF genes and correlates with H3K9me3 enrichment in K562 cells.

### ATRX enrichment at ZNFs is conserved across distinct human cell-types

Next, we investigated whether ATRX is bound at ZNF 3’ exons in other cell types. Because ATRX, SETDB1 and TRIM28 ChIP-seq data sets are not readily available, we first utilized available ZNF274 and H3K9me3 ChIP-seq data in ENCODE for cell lines GM12878, H1-hESC, HeLa-S3, HepG2 and NT2-D1 (see Table S6 for details of data sources and analysis). ZNF274 binds to the 3’ of ATRX-bound Class I and Class II ZNF genes (as defined in K562) in the majority of cell lines analyzed (**Fig. S4C**). Similar to other previous studies, ^44^ H3K9me3 at the 3’ region of ZNFs was also conserved (**Fig. S4C**).

Using a panel of cell lines of diverse origin from ENCODE (K562, H1-hESC, HeLa-S3 and HepG2) and two neuroblastoma cell lines (LAN6 and SKNFI), we performed ATRX ChIP-qPCR for eleven randomly chosen Class I ZNFs distributed across different chromosomes, as well as two Class III ZNF genes as negative controls. We found significant enrichment of ATRX in Class I ZNFs across all cell lines analyzed, and none for Class III ZNFs or IgG control (**Fig. 3A**). <u>Using available ChIP-seq data, we plotted the enrichment of H3K9me3 and ZNF274 at these specific ZNFs in the ENCODE cell line panel</u>. As expected, all ATRX-enriched ZNFs also show relatively high levels of H3K9me3 and ZNF274 (**Fig. 3B**). Furthermore, <u>these ZNFs show diverse expression levels independent of their ATRX binding status according to the normalized RNA-seq signal from ENCODE (**Fig. 3C**)</u>. This result substantiates the notion that ATRX binding is independent of the transcriptional status of the ZNFs.

**Figure 3.**
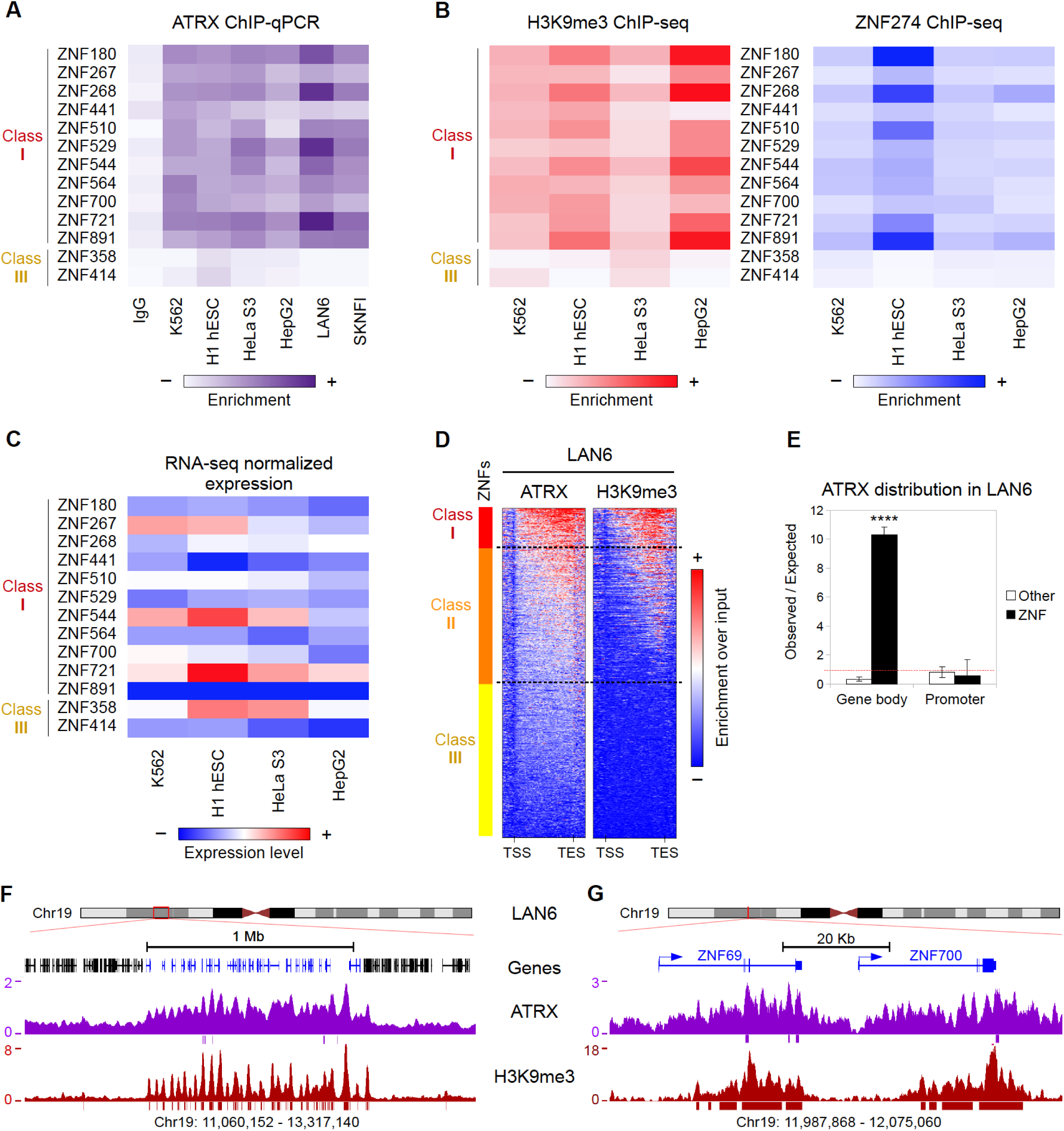
ATRX binding to ZNF 3’ exons is conserved across human cell lines. (A) Heatmap of ATRX ChIP-qPCR enrichment over ZNF genes in several human cell lines. The color represents the average enrichment of at least 2 independent biological replicates per cell line. ZNF class I genes show significant enrichment as compared to IgG for all assessed cell lines (for details see **Table S3**). (B) H3K9me3 (left) and ZNF274 (right) ChIP-seq enrichment over the same panel of ZNFs as in (A) for cell lines with available ENCODE data. (C) RNA-seq normalized expression values (log2(RPKM*1)) from ENCODE for the panel of ZNF genes shown in (A) and (B). (D) Metagene profiles of ATRX and H3K9me3 ChIP-seq data over ZNFs gene bodies +/- 1kb in LAN6 neuroblastoma cell line. (E) Observed over expected random distribution of significant ATRX peaks in ZNFs and non-ZNF (other) promoters and gene bodies for LAN6. (F) UCSC Genome browser screenshot of a typical ZNF cluster (genes in blue) on chromosome 19. The enrichment over input signal for ATRX and H3K9me3 ChIP-seq in LAN6 is shown. Significant peaks represented as bars below enrichment tracks. (G) Zoomed in snapshot of two ZNFs contained within the ZNF cluster shown in (F).

In order to corroborate our findings genome-wide, we performed ATRX and H3K9me3 ChIP-seq in LAN6, which showed robust ATRX enrichment by ChIP-qPCR (**Fig. 3A**). Strikingly, we found that the binding of both ATRX and H3K9me3 at the 3’ of ZNF genes is conserved in LAN6 (**Fig. 3D**, compare with **Fig. 2D**). This binding was specific and highly significant for ZNF gene bodies (**Fig. 3E**), and as observed in K562 cells, ATRX and H3K9me3 demarcate the ZNF clusters on chromosome 19 in LAN6 (**Fig. 3F–G**).

Next, we re-analyzed the only other published human ATRX ChIP-seq dataset from primary human erythroblasts ^15^ and two recent datasets from mouse embryonic stem cells (mESC) and mouse embryonic fibroblasts (MEF), ^46^ and assessed ATRX binding at ZNFs. In contrast to K562, LAN6 and all other cell lines analyzed, human erythroblasts, mESCs and MEFs lack ATRX binding within ZNF gene bodies (**Fig. S4D,E**). We then reanalyzed available ChIP-seq data for H3K9me3 in mESCs and MEFs (Table S6) and found that H3K9me3 enrichment at ZNFs does not co-localize with ATRX, and only mESC showed H3K9me3 enrichment at the 3’ exons of ZNFs (**Fig. S4E**). It is unclear if these differences are biologically significant and it will be interesting to define if the pattern observed in humans is also conserved in other vertebrates.

Overall, our analysis demonstrates that the enrichment pattern of ATRX, H3K9me3 and ZNF274 at ZNF 3’ exons is conserved in several human cell lines of diverse lineages, including non-tumorigenic cells, suggesting a general mechanism for ZNF gene regulation via ATRX that is independent of transcription.

### ATRX deficient cells have reduced H3K9me3 enrichment at 3’ exons of ZNFs

To functionally investigate the role of ATRX, we generated two clonal ATRX knock out (KO) cell lines using CRISPR/Cas9 genome editing in K562 cells. Our KO lines (ATRX KO1 and KO2) were characterized in detail (**Fig. S5A**, Methods). As a control, we used a clonal cell line overexpressing Cas9 alone (referred to as V2). Immunoblot and sequencing analyses showed that ATRX KO1 cells retain residual ATRX from an allele with an in-frame deletion, while KO2 cells are completely devoid of ATRX protein (**Fig. 4A, S5A**). <u>Moreover, the global levels of SETDB1, TRIM28 and ZNF274 in chromatin were largely unaffected by ATRX KO (**Fig. S5B**).</u> We next performed ATRX ChIP-qPCR in these KO lines for the panel of Class I and Class III ZNFs. In accordance with our ChIP-seq data, Class III ZNFs lacked ATRX enrichment, while Class I ZNFs were enriched for ATRX in the control, reduced in KO1 and ablated in KO2 cells (**Fig. 4B**).

**Figure 4.**
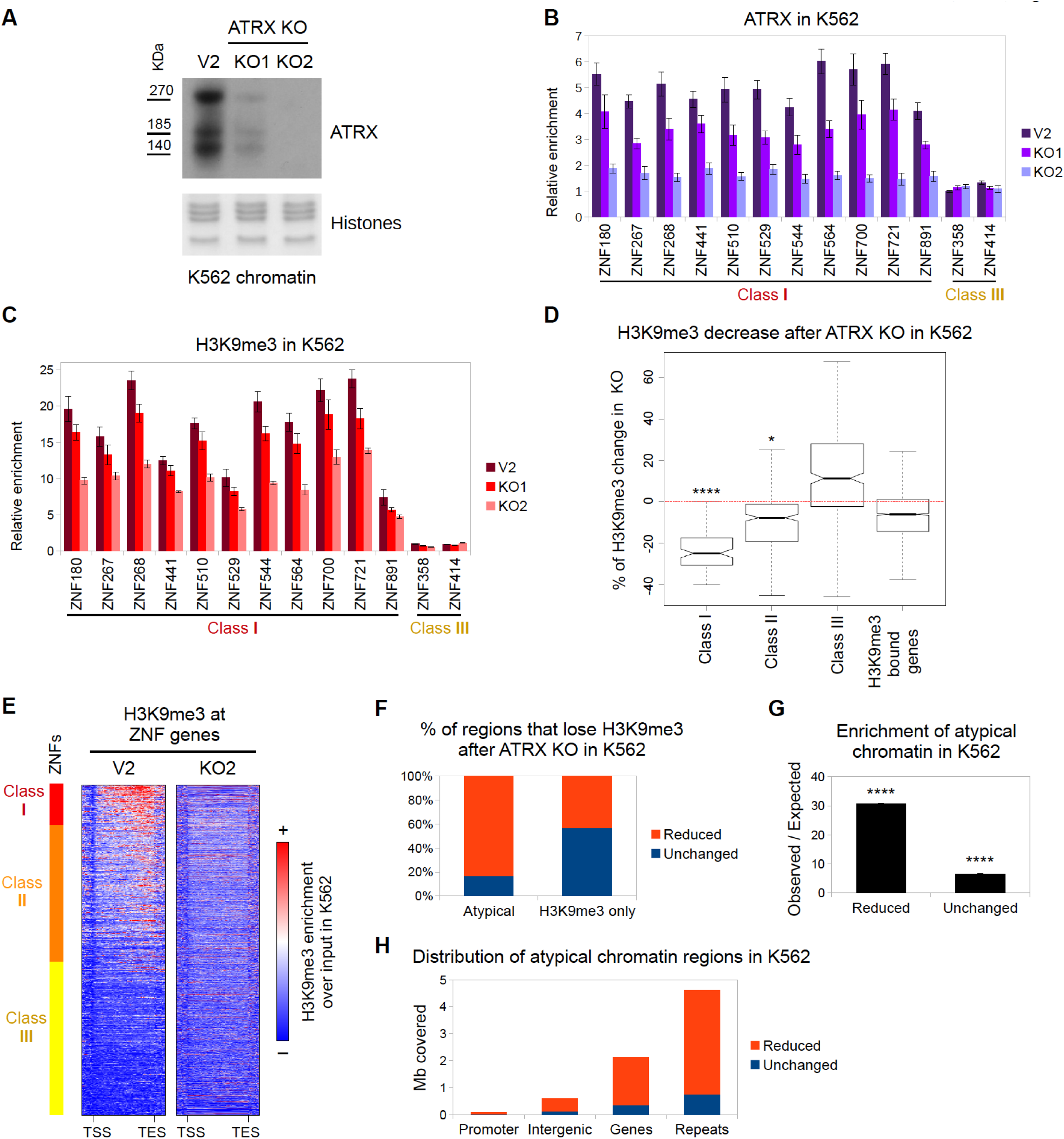
ATRX deficient cells have decreased levels of H3K9me3 at 3’ exons of ZNFs and other atypical chromatin regions. (A) Western blot of ATRX in chromatin preparations from control (V2) and two CRISPR ATRX KO K562 cell lines. Amido Black staining of histones is shown as a loading control. (B) ATRX ChIP-qPCR over ZNF genes in control and ATRX KO K562 cell lines. (C) Same as in (B) with H3K9me3 native ChIP-qPCR. In both (B) and (C), bars represent average of at least 3 biological replicates and the error bars represent the SEM. (D) Box plot of the H3K9me3 changes (represented in % with respect to the control) observed in the ZNF classes and non-ZNF H3K9me3-bound genes upon ATRX KO. Asterisks show significant changes with respect to the non-ZNF genes (* pvalue < 0.05; **** pvalue < 1×10^−4^). (E) ChIP-seq metagene profile of H3K9me3 in control (V2) and ATRX KO (KO2) K562 cell lines over ZNFs. (F) Distribution of reduced and unchanged H3K9me3 regions after ATRX KO in atypical chromatin and H3K9me3-only regions. (G) Observed over expected distribution of reduced and unchanged regions at atypical chromatin. Asterisks show significantly overrepresented regions (pvalue < 1×10^−4^) (H) Genomic distribution of atypical chromatin regions.

Loss of ATRX has been shown to promote genomic instability and defects in cell cycle. ^37–40^ We tested these functional readouts in our KO cell lines and found that ATRX KO K562 cells have increased DNA damage as compared to control cells as assessed by Comet assays and immunoblots for γH2A.X (**Fig. S6A-C**). Furthermore, ATRX KO cells displayed a slight, but reproducible, cell cycle defect in the G1/S transition (**Fig. S6D,E**).

We then queried whether ATRX deficiency alters the chromatin state of the atypical chromatin domains at ZNF genes. To address this question, we performed native ChIP for H3K9me3 and H3K36me3 in control and ATRX KO cells, followed by qPCR analysis for ZNFs. Strikingly, H3K9me3 levels decreased at Class I ZNFs in the ATRX KO cells compared to control cells, and displayed a similar pattern to that of ATRX occupancy (**Fig. 4B, C,**). In contrast, H3K36me3 levels remain stable upon ATRX loss (**Fig. S7A**). We also assessed whether ATRX depletion affected the transcription of ZNF genes. Consistent with our findings that ATRX enrichment at ZNFs does not correlate with their transcriptional activity (**Fig. 2C, 3C**), we did not observe changes in expression levels of ZNFs in the ATRX KO cell lines (**Fig. S7B**). Together, these data suggest that ATRX regulates H3K9me3 levels at ATRX-bound ZNF genes, but that H3K36me3 and transcriptional regulation is ATRX-independent.

Because ZNF274 binds DNA and recruits TRIM28, which in turn recruits SETDB1, we speculated that the ZNF274/TRIM28/SETDB1 complex serves as a scaffold for ATRX at the 3’ exons of ZNFs. Therefore, we posited that ATRX KO would not affect the complex binding to the chromatin, but rather its capacity to deposit or maintain H3K9me3. As expected, ZNF274 and TRIM28 binding remains largely unchanged at ZNF genes after ATRX KO (**Fig. S7C,D**). While ATRX KO1 cell line did not affect SETDB1 binding, SETDB1 occupancy was increased in ATRX KO2 (**Fig. S7E**). This may reflect a compensatory effect for the complete loss of ATRX, although we cannot rule out a clone-specific effect. These data broadly favors a model in which ZNF274/TRIM28/SETDB1 is able to bind to the 3’ exons of ZNFs independently of ATRX, but ATRX is required to establish or maintain H3K9me3 at atypical chromatin of ZNF genes.

To investigate the genome-wide alterations of H3K9me3 after ATRX depletion, we performed native ChIP-seq for H3K9me3 in control and ATRX KO cells. We first analyzed the H3K9me3 enrichment at ZNF genes. In agreement with our ChIP-qPCR results, we found H3K9me3 levels significantly reduced in KO cells, specifically in ATRX-enriched Class I and II ZNFs (**Fig. 4D**). These results were confirmed by metagene analysis of ZNFs (**Fig. 4E**) and visualization of H3K9me3 peaks at the ZNF clusters on chromosome 19 (**Fig. S8A, B**). This demonstrates that ATRX is required for establishing and/or maintaining H3K9me3 levels at the 3’ end of a subset of ZNFs.

H3K9me3 within atypical chromatin domains is sensitive to ATRX, but not DAXX, depletion Besides ZNF genes, we identified additional genomic regions that show reduction of H3K9me3 upon ATRX depletion (**Fig. S8C**). To further understand the nature of the H3K9me3 regions that are sensitive to ATRX loss, we divided all H3K9me3 regions to “reduced” (n=6,055) or “unchanged” (n=8,331) based on whether the levels are decreased or remain stable in the ATRX KO cell line, respectively. We then analyzed the proportion of reduced regions in atypical chromatin and compared it to H3K9me3-only chromatin. Strikingly, a large proportion of reduced regions (~80%) were atypical chromatin in contrast to H3K9me3-only chromatin (~40%) (**Fig. 4F**). This trend is highly significant (**Fig. 4G**). Thus, our data suggest that the levels of H3K9me3 at atypical chromatin are particularly sensitive to ATRX depletion.

We next analyzed the genomic distribution of atypical chromatin. Interestingly, as shown in **Figure 4H**, most atypical chromatin regions that lose H3K9me3 are located in repetitive sequences, followed by those located in genes (including ZNF genes). This is consistent with recent studies that show a role for ATRX in regulating H3K9me3 at repetitive elements, however the levels of H3K36me3 at such regions have not been described. ^7,20–22,25^

ATRX has been reported to regulate H3K9me3 levels in concert with DAXX at a subset of repetitive and imprinted regions. ^7,20–22,25^ In order to assess whether DAXX regulates H3K9me3 at ZNFs, we used CRISPR/Cas9 to generate DAXX KD and double DAXX KD * ATRX KO lines (see Methods for details). After confirming the knock down of DAXX (**Fig. 9A**), we performed H3K9me3 ChIP-qPCR for Class I and Class III ZNFs. We found little to no change in H3K9me3 levels in the single DAXX KD (**Fig. 9B**). The DAXX KD/ATRX KO shows a consistent decrease of H3K9me3 as observed with the single ATRX KO2 cell line (**Fig. 9B**). While we can not completely rule out a role for DAXX in the maintenance of H3K9me3 levels at ZNFs, our results suggests that its contribution is minor in comparison to other factors, such as ATRX and ZNF274 (see below). Overall, these results suggest that ATRX protects H3K9me3 levels at atypical chromatin regions genome-wide, and that ZNF genes comprise a subset of these affected regions.

### ZNF274 recruits ATRX to a subset of ZNF genes to regulate H3K9me3 levels and genomic stability

To further understand the interplay between ATRX and ZNF274, we depleted ZNF274 using CRISPR/Cas9 <u>in K562 cells</u>. We generated three clonal cell lines (see Methods for details): a single ZNF274 KO cell line (referred to as Z274 KO), a double ZNF274/ATRX KO cell line (Z274/ATRX KO) and a control cell line expressing a non-specific sgRNA (Random). We characterized the mutation status of the ZNF274 gene (**Fig. S10**) and confirmed that the Z274 KO abrogated ZNF274 binding to ZNF genes by ChIP-qPCR (**Fig. 5A**).

**Figure 5.**
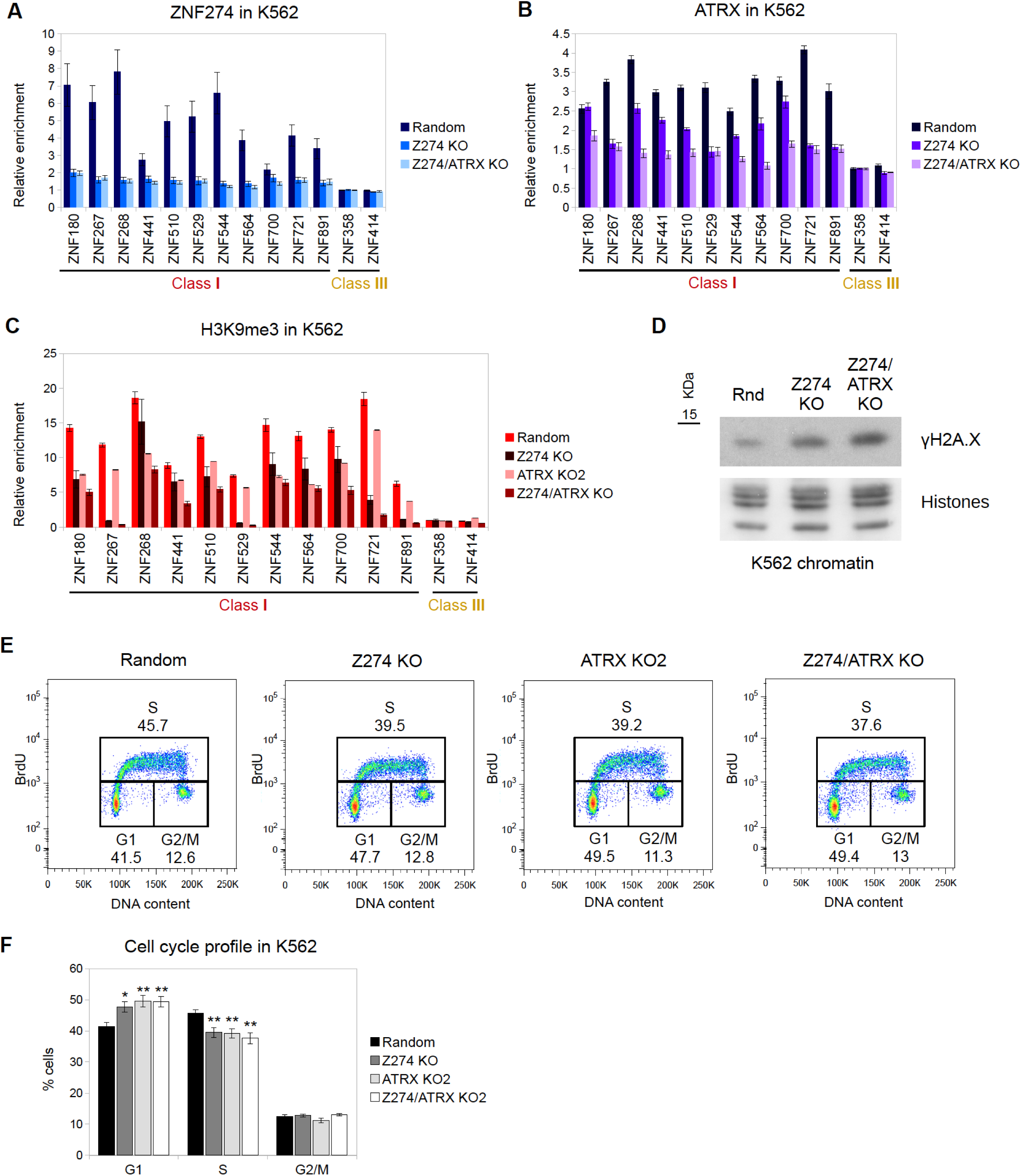
ZNF274 KO reduces ATRX and H3K9me3 levels at ZNFs. (A) ZNF274, (B) ATRX and (C) H3K9me3 ChIP-qPCR in <u>K562</u> ZNF274 KO and <u>K562</u> double ZNF274/ATRX KO at ZNF genes. Single ATRX KO2 cells are used for the H3K9me3 ChIP for comparison. In all graphs, the bars represent the average of at least two independent biological replicates. Error bars depict SEM. Results of statistical comparisons in **Table S3**. A non-specific sgRNA (random) used as control (see **Table S7**). (D) <u>Chromatin</u> immunoblot of yH2A.X in control (Rnd) and ZNF KO <u>K562</u> cells. Histones used as loading control. (E) Representative <u>K562</u> cell cycle profiles of control (random), ATRX and ZNF274 single and double KO assessed by BrdU/PI staining. n > 6 biological replicates. (F) Graph depicting quantifications of (E). The bars show the average % of cells in each phase, error bars depict SEM. Asterisks show significant changes compared to the control (* pvalue < 0.05; ** pvalue < 0.01).

We next analyzed ATRX binding in the ZNF274 KOs by ChIP-qPCR. Strikingly, we observed a decreased binding of ATRX in the Z274 KO cells for most Class I ZNFs. As expected, ATRX is depleted from all ZNFs in the Z274/ATRX KO (**Fig. 5B**). <u>These results strongly suggest that ZNF274 tethers ATRX to ZNFs. However, because ATRX is not lost at all ZNFs examined, we hypothesize that there may be other (potentially compensating) factors that can recruit ATRX to these regions</u>.

To determine the impact of ZNF274 depletion on H3K9me3, we performed ChIP-qPCR in the ZNF274 KO lines. In order to compare the effect of the double Z274/ATRX mutant, we included the ATRX KO2 cell line in our analysis. <u>As shown in **Figure 5C**, the ZNFs that have lost ATRX binding in the ZNF274 KO cells also lose H3K9me3 enrichment. In contrast, the ZNFs that still retain some ATRX binding show a more modest reduction in H3K9me3 levels.</u> Strikingly, in many cases, the Z274/ATRX KO cells showed a stronger decrease of H3K9me3 than either individual KO, suggesting that knocking out a single component of the complex may not entirely abrogate its function.

Finally, we determined the functional consequence of H3K9me3 reduction in the ZNF274 KOs. We noted an increase in γH2A.X in the Z274 KO cell line as compared to the control, which was even stronger in the Z274/ATRX KO cells (**Fig. 5D**). Because the ATRX KO cells show a defect in cell cycle, we queried whether the ZNF274 KO would have a similar phenotype. Indeed, ZNF274 KO cells show a delay in G1/S (**Fig. 5E–F**), comparable to that observed in the ATRX KO2 cells.

Collectively, we propose a model in which ATRX is tethered to the 3’ exons of ZNF genes by the ZNF274/TRIM28/SETDB1 complex to establish or maintain/protect H3K9me3 at these transcriptionally active regions (**Fig. 6**). ATRX and ZNF274 depletion leads to reduction of H3K9me3 levels at ZNF 3’ exons and is associated with higher DNA damage and cell cycle defects. Furthermore, ATRX acts as a genome-wide regulator of H3K9me3 levels at atypical chromatin that is not restricted to ZNFs. Finally, we propose that impairing ATRX function has important consequences for the genomic stability and evolution of ZNFs clusters, as the loss of H3K9me3 at ZNF 3’ exons may increase the probability of HR between them (**Fig. 6**; see discussion below).

**Figure 6.**
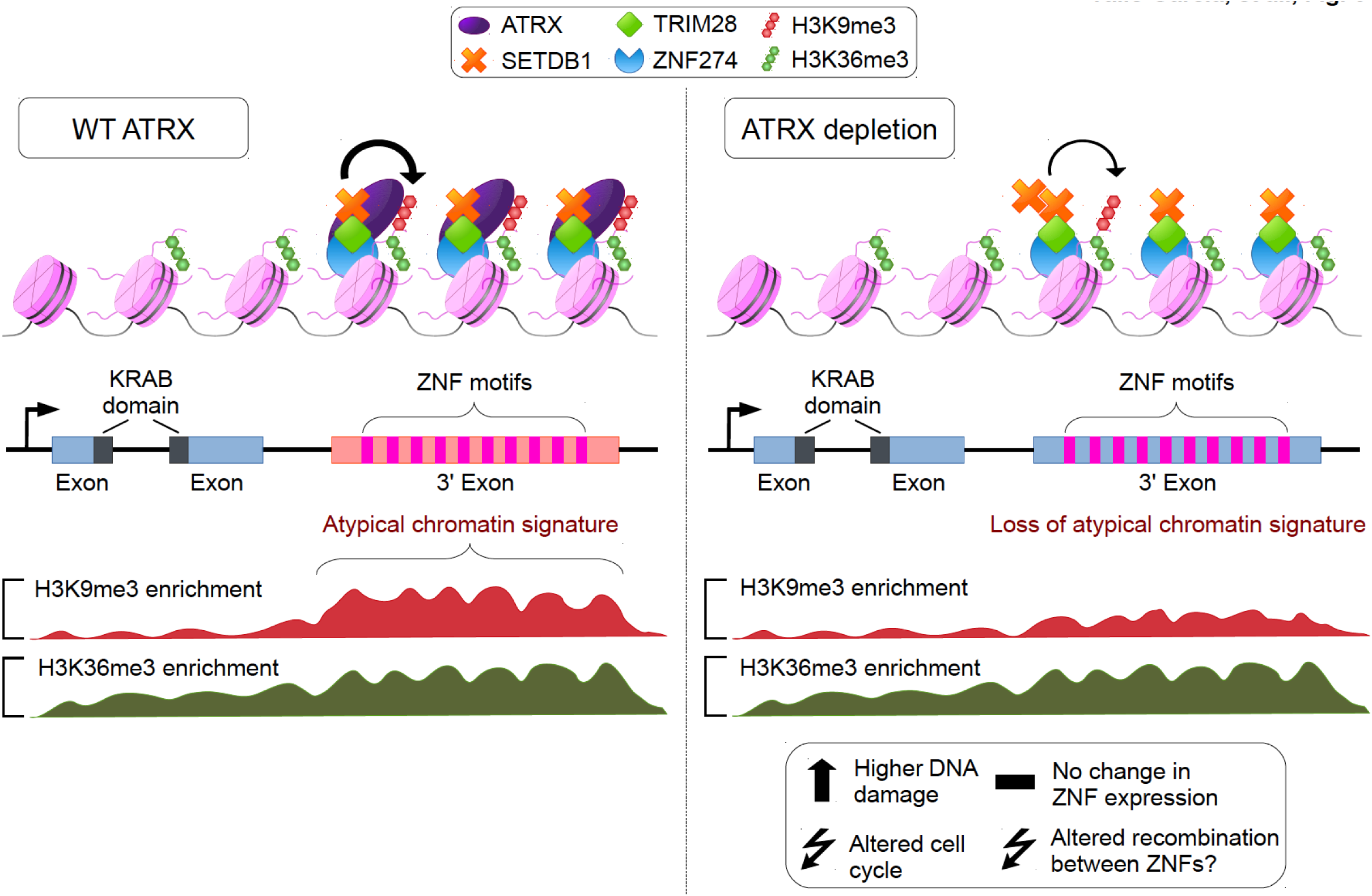
Model of ATRX regulation at ZNF 3’ exons. Left: <u>ATRX forms a complex with ZNF274, TRIM28 and SETDB1 to facilitate the deposition and maintenance of H3K9me3 at ZNF 3’ exons.</u> The presence of the mark establishes an atypical H3K9me3/H3K36me3 domain. Right: <u>Upon ATRX depletion</u>, H3K9me3 and the atypical chromatin domains at ZNF 3’ exons are lost. Loss of ATRX induces altered cell cycle, increased DNA damage and possibly recombination between ZNFs.

## DISCUSSION

ATRX is an important chromatin regulator involved in diverse processes such as transcriptional regulation, maintenance of imprinted loci, replication, genome stability and chromatin looping. ^7–9,30,54^ Here we report that ATRX binds the 3’ exons of ZNFs with an atypical chromatin signature to establish or maintain H3K9me3 (**Fig. 6**). ZNFs represent the fastest expanding gene family encoding transcription factors in humans, with the greatest diversity of target sequences. ^41–43,55^ This property of ZNFs to recognize a multitude of motifs is generated through diverse combinations of their zinc finger motifs. ^55^ The average ZNF gene contains 9 independent zinc finger motifs and the number of motifs varies from gene to gene. Interestingly, these motifs are almost always located at the 3’ exons of ZNF gene loci and harbor an atypical chromatin signature consisting of both H3K9me3 and H3K36me3 (**Fig. 6**). ^42,44^

Through an unbiased ChIP-seq approach and analysis of ENCODE data, we found that ATRX preferentially binds to the 3’ exons of a subset of ZNF genes containing this atypical chromatin signature. These ZNF genes are distinguished by the presence of a KRAB repressor domain, a higher than average number of zinc finger motifs, low levels of G-content and low probability of forming G-quadruplexes. Previous studies have shown that ATRX recognizes and resolves G-quadruplexes, relevant in the context of gene regulation. ^15^ However, our data suggests that ATRX binds and regulates these atypical chromatin regions by an alternative mechanism(s).

We found that ATRX co-localizes and interacts with the previously reported ZNF274/TRIM28/SETDB1 complex ^53^ at ZNF 3’ exons with an atypical chromatin signature. Our data suggests a model whereby ZNF274 recognizes a DNA motif that is common in certain ZNF 3’ exons (i.e. those bound by ATRX), <u>and recruits the TRIM28/SETDB1/ATRX complex</u> to establish, maintain or protect H3K9me3. <u>Of course, there are questions about this model that require further clarification.</u> For example, we wonder why in the absence of ZNF274, ATRX still retains its ability to bind a subset of ZNFs. These results, coupled to the fact that we find atypical chromatin regions that are enriched in TRIM28/SETDB1/ATRX but not ZNF274 (data not shown), suggests that other ZNF transcription factors may be involved in targeting ATRX to specific loci. Thus, the TRIM28/SETDB1/ATRX complex may be guided to unique loci or chromatin environments via ZNF proteins that may have cell- or stage-specific binding patterns. Moreover, while it has been established that KRAB-ZNF proteins recruit TRIM28 through their KRAB domain, and that KRAB in turn recruits SETDB1 ^56^, the dynamics of ATRX recruitment to the complex are unclea<u>r. One possibility is that SETDB1 establishes basal H3K9me3 levels that are recognized and bound by the ATRX ADD domain, which in turn promotes further H3K9 methylation forming a feedback loop to maintain H3K9me3 levels</u>.

Our loss-of-function studies revealed that H3K9me3 levels at atypical chromatin regions, in particular those at ZNF 3’ exons, are decreased after ATRX loss. Recent studies have reported similar effects upon ATRX loss at known ATRX target regions such as ERVs and imprinted loci. ^20–22,25^ However, the mechanisms underlying H3K9me3 loss in ATRX-deficient cells remain unclear. Based on our observations, we hypothesize three nonexclusive scenarios: 1) ATRX may facilitate SETDB1-mediated H3K9me3 deposition by promoting an optimal nucleosome structure for SETDB1 function, 2) ATRX binds H3K9me3 and blocks demethylase activity, and 3) ATRX may help to reestablish H3K9me3 after transcription or replication. These scenarios would also explain why H3K9me3 levels decrease after ATRX KO, despite the fact that SETDB1 binding remains unchanged.

Functionally, loss of H3K9me3 in ATRX and ZNF274 KO cells may impact on the genomic stability of ZNFs. The ZNF family has expanded in the primate lineage through gene duplication. ^41–43^ ZNFs contain long stretches of highly similar sequences in their 3’ exons and high levels of H3K36me3, a mark that enhances the resolution of double strand breaks (DSB) through the HR pathway. ^57,58^ Therefore, ZNF regions may be prone to recombine. As H3K9me3-rich heterochromatin regions are refractory to repair of DSB and HR, ^59,60^ we hypothesize that ATRX-mediated H3K9me3 enrichment at ZNF 3’ exons protects them from HR. In this regard, recent studies demonstrated that ATRX can act as a suppressor of recombination at telomeres. ^22,36,47,61^ Furthermore, the notion that H3K9me3 at ZNF 3’ regions protects from HR has been previously suggested in at least two independent studies. ^44,49^ If this hypothesis is correct, ATRX may function, at least partially, as an epigenetic modulator of ZNF gene expansion, balancing the positive selective pressure for ZNF duplication and the deleterious effects that an excessive recombination rate has for genomic stability. Moreover, altered recombination levels at ZNFs, telomeres and other ATRX-regulated regions may be responsible, at least in part, for the defects in cell cycle and an increased percentage of DNA damage we observed in ATRX KO cells.

Despite the evidence that ATRX suppresses recombination at certain genomic regions, there are alternative explanations for its role at ZNFs. For instance, we find that Class I ZNFs tend to be late replicating. Thus, it is possible that ATRX facilitates DNA replication at atypical chromatin regions. This idea is in line with the finding that ATRX KO cells subjected to replicative stress have a higher accumulation of stalled replication forks. ^39,40^ Moreover, a replication defect can also explain the higher levels of DNA damage and the cell cycle delay observed in ATRX and ZNF274 KO cells. Furthermore, it has been suggested that the chromatin environment can affect RNA polymerase processivity, in turn affecting the RNA splicing machinery. ^62^ Thus, the levels of H3K9me3 at ZNFs could affect alternative splicing. Because the RNA-seq analyses performed in this study were focused only on global expression levels, potential changes in isoform expression, polymerase processivity, and other related mechanisms, remain to be explored.

Several recent studies have found that the ATRX/DAXX complex deposits the H3.3 histone variant at several H3K9me3-containing regions, such as telomeres, imprinted regions and retrotransposons. ^7,2–22,25^ Interestingly, H3K9me3 levels in those regions are frequently decreased upon ATRX inactivation. How similar these H3.3/H3K9me3 regions are to the H3K9me3/H3K36me3 atypical chromatin signature remains an open question. While our data suggests that DAXX has a minimal role in maintaining H3K9me3 at ZNF 3’ regions, we cannot exclude the possible involvement of H3.3.

In summary, we demonstrate here that ATRX regulates H3K9me3 levels at the 3’ exons of ZNFs and other loci containing an atypical chromatin signature. This unexpected function sheds light onto the complex genomic regulatory pathways that ATRX participates in, and may be important for the future understanding of diseases in which ATRX is mutated or altered.

## MATHERIALS AND METHODS

### XL-MNase ChIP

Cross-linked-MNase ChIP was performed with Cell Signaling SimpleChIP Enzymatic Chromatin IP Kit (cat. #9003) following manufacturer’s instructions with modifications. Briefly, K562 and LAN6 (2-4 × 10^6^) cells were cross-linked with 1% formaldehyde in PBS, 100mM NaCl, 1mM EDTA pH 8.0, 50mM HEPES pH 8.0, for 10 minutes at room temperature. Reaction was quenched with 125 mM glycine. Cells were lysed to obtain nuclei and chromatin was digested with Micrococal Nuclease (MNase) (NEB, cat. #M0247S) at 37°C for 20 minutes. Nuclei were disrupted by brief sonication (4 cycles, 20 sec ON/OFF, high power) in a Bioruptor Twin. Chromatin was quantified and 40-80 ug was incubated with antibody at 4°C overnight with 1% taken as input sample. After incubation, Magna protein A/G magnetic beads (Millipore, cat. #16-663) were added for 3 hours at 4°C. Beads were washed following the manufacturer’s protocol, followed by an extra LiCl buffer wash (10mM Tris-HCl pH 8.0, 1mM EDTA pH 8.0, 1% sodium deoxicholate, 1% Igepal, 250mM LiCl). DNA bound to complexes was eluted at 65°C for 30 mins, treated with RNAse A (10mg/ml) for 1 hour at 37°C, with Proteinase K (20mg/ml) for 3 hours at 55°C and then cross-linking was reversed for 4-6 hours at 65°C. DNA was purified using the Qiagen MinElute PCR purification kit and subsequently analyzed and quantified using an Agilent 2100 Bioananalyzer High Sensitivity Kit.

### Native ChIP

Native ChIP was performed as previously described ^63^ with minor changes. Briefly, nuclei isolated from K562 and LAN6 (4-6 × 10^6^) cells were treated with MNase (Affymetrix Cat #70196Y) and ~40-80ug of digested chromatin was immunoprecipitated with specific antibodies. The immunoprecipitated material was treated with Proteinase K for 3h at 56 °C and purified using the Qiagen MinElute PCR purification kit.

### Antibodies

See **Table S4** for a full list of antibodies and concentrations used for each assay.

### Library preparation and ChIP-seq

ChIP-seq libraries were prepared as previously described ^63^ and libraries were sequenced single-end on Illumina HiSeq 2500 and NextSeq 500. See **Table S5** for detailed descriptions (number and size of reads, etc.) of the sequenced samples.

### ChIP-seq analysis

Sequenced reads were aligned to the GRCh37 (hg19) assembly using Bowtie 1.0.0. ^64^ Redundant reads were eliminated using MACS2 (2.1.0) ^65^ filterdup option with default parameters. The estimated background reads and optimal normalization factor between ChIP and Input samples was calculated with the R NCIS package 1.0.1 ^66^ using a shift size of 75bp. Peak calling was performed with MACS2 callpeak using the --ratio option with the estimated value from NCIS. ChIP/Input fold enrichment pileups were created with the macs2 bdgcmp tool using the -m FE option and converted to bigWig files using the bedGraphToBigWig program (v4) from the UCSC binaries. For some samples, a second peak calling was performed using SICER ^67^ 1.1 using a window of 200bp, allowing gaps of 400pb and filtering for q-values < 1×10^−8^ (see **Table S6** for details). All peaks overlapping with ENCODE blacklisted regions ^45^ were eliminated. When available, a final list of peaks overlapping the MACS2 and SICER peaks was obtained from the intersectBed program from bedtools 2.17.0. ^68^ See **Table S6** for GEO accessions, details of analysis parameters (q-values, etc.), and results for all the datasets analyzed.

For all the ATRX-related analysis, the K562 ATRX-H300 ChIP-seq sample was used due to its high quality, low percentage of background, and high number of reads (**Table S5**).

### Gene and coordinate analyses

All analyses were performed using Ensembl genes (putative genes, pseudogenes and genes unmapped to chromosomes were excluded). For human the Ensembl genes 75 version (GRCh37.p13) was used. For mouse the Ensembl genes 77 version (GRCm38.p3) was used. The lists of ZNF genes were downloaded from Biomart, using the InterPro id for the Zinc Finger C2H2 domain (IPR007087) as a filter, manually analyzed and curated. Repeats coordinates were obtained from the repmask table from the UCSC Genome Browser for hg19. Bed files for subsequent analysis were generated using the following coordinates: promoters (-3kb to TSS), intergenic (regions falling outside genes or promoters).

### Correlation analysis

Correlation heatmaps were generated with the bigwigCorrelate program from the DeepTools suite (v 1.5.9.1) ^69^ using the spearman correlation method. Fold enrichment over input bigwig files were used as inputs. For genome-wide analysis, all chromosomes were divided in 10kb non-overlapping bins. Bins falling in the blacklisted regions from Encode were excluded. For gene-specific analysis, a bed file with the coordinates from TSS to TES was used.

### Analysis of genomic distribution and correlation with chromatin states

Bed files with the coordinates of the chromatin states for K562 calculated in ^48^ were generated. Similar chromatin states were merged into single categories (states 1-3 were fused as promoters, states 4-7 as enhancers, 10-11 as transcription, 14-15 as repetitive). The probability of overlap between ATRX peaks and the HMM states, genes, promoters and intergenic regions was calculated with The Genomic HyperBrowser. ^70^ ATRX peaks were randomized 10,000 times preserving its segment length and intersegment gaps. Observed/expected values were calculated by dividing the overlap of the ATRX peaks over the overlap of the randomized regions.

### Gene Ontology analysis

Genes overlapping ATRX significant peaks were obtained using intersectBed from the Bedtools suite. Gene Ontology analysis was performed with DAVID ^71^ using default parameters. See **Table S1** for the list of genes overlapping ATRX peaks.

### Generation of ZNF classes

Average ATRX ChIP/Input enrichment per gene was obtained with the computeMatrix program from the DeepTools suite. The genes were then clustered by kmeans into 3 groups using R (v 3.0.1). Bed files from each ZNF Class are provided in **Additional File 1**.

### Protein domain analysis

The protein sequence of the ZNFs was obtained with Biomart. The sequences were matched to all the Prosite motifs using the ScanProsite tool with default parameters. Inhouse scripts were used to parse the ScanProsite output and count the number of motifs per ZNF gene.

### G-content and g-quadruplex analysis

G-content at the 3’ region of ZNF genes (last 3kb) was calculated with in-house scripts. The probability of G-quadruplex formation was calculated using the quadparser program ^72^ with default parameters.

### Average enrichment analyses

The color plots showing the distribution of genetic features among ZNF classes were drawn using the image function of R. Darker colors represent presence of KRAB domains, a higher number of zinc finger motifs, high G content at the C-terminal ZNF region (last 3kb of the gene) and presence of sequences predicted to form Gquadruplexes. For RNA-seq data, the Z-score of the RNA-seq normalized signal (log2(RPKM*1)) from K562 ENCODE data was calculated and plotted. Red colors match high expression signals, while blue colors match low expression signals. ZNFs are sorted from high to low ATRX enrichment from top to bottom. The calculated values used to generate color plots of **Figure 2C** are provided in **Additional File 2**.

### Metagene analysis

ZNF genes were ordered by ATRX enrichment from high to low. The metagene plots were generated with DeepTools using the ordered gene file and the ChIP/Input bigwig tracks. All genes from TSS to TES were scaled to a 5kb region +/-1kb with sliding windows of 100 bp. Metagene values were calculated using computeMatrix. Values were plotted with heatmapper and the average enrichment profiles were plotted with profiler. The matrices for all heatmaps and profile plots are provided in **Additional File 3**. The data sources can be found in **Table S6**.

### Motif analysis

The ZNF274-bound regions were obtained from the summit peaks generated by MACS2. A fasta file from these coordinates was created using in-house scripts and used as input for MEME-ChIP (v. 4.1.0.0). ^73^ The sequences were compared to the JASPAR vertebrate database; other options were set as defaults. The regions matching the motifs (fimo output) were parsed, sorted by class and counted using in-house scripts.

### Repli-seq analysis

Repli-seq bigwig files generated by the ENCODE project for K562 were analyzed using DeepTools. The average signal per ZNF was calculated with computeMatrix and then plotted in R using the heatmap.2 function. Calculated Repli-seq values are provided in **Additional File 4**.

### Co-immunoprecipitation of chromatin bound proteins

K562 cells were harvested for each IP (50 × 10^6^) using PBS supplemented with protease inhibitor without EDTA, 1 mM DTT and 0.05% NP-40 and incubated on ice for 3 min and centrifuged at 500 × g for 5 min at 4°C. The nuclei were washed with PBS and resuspended in EX 100 buffer (10 mM Hepes pH 7.6, 100 mM NaCl, 1.5 mM MgCl2, 0.5 mM EGTA, 10% glycerol) and digested with MNase (NEB, cat. #M0247S) at 37°C for 10 min. The reaction was stopped with EGTA (10mM) followed by spinning (10,000xg for 10 min at 4°C). The collected supernatant containing mononucleosome particles was used for IP. 4μg of ATRX antibody (HPA001906) or Rabbit IgG were added to each IP and incubated overnight (rotating at 4°C). 2 μl of protein A*G magnetic bead slurry (Millipore, cat. #16-663) were washed twice with PBS and once with EX 100 buffer and incubated with the immunocomplexes for 2.5 hours at 4°C followed by 2 × 5 min washes with buffer G250 (50 mM Tris pH 7.5, 250 mM NaCl and 0.5% NP-40) and one 5 min wash with G150 (50 mM Tris pH 7.5, 150 mM NaCl and 0.5% NP-40). Laemmli sample buffer was added to each sample, boiled at 95°C, and run on NuPAGE 4-12% bis-tris gels (Thermo Fisher).

### Generation of ATRX and ZNF274 KO cell lines by CRISP/Cas9

Single guide RNAs (sgRNAs) targeting the exons of the ATRX gene were designed ^74^ and cloned into the lentiCRISPR V2 vector (addgene #52961, see sgRNAs sequences in **Table S7**). Lentiviral production using HEK293T cells was performed using standard laboratory protocols. To generate stable cell lines, K562 (~1x10^6^) cells were infected with virus for each CRISPR guide and either an empty lentiCRISPR V2 or a lentiCRISPR V2 containing a non-specific sgRNA was used as a control. Infected cells were selected (puromycin 2ug/ml; 3 weeks) and subsequently single clones were sorted into 96-well plates using an IMI5L cell sorter (BD Biosciences) for ATRX and ZNF274 KO. Clones were grown in selection for 3-4 weeks and tested for KO by DNA sequencing and immunoblot analysis. For the generation of ATRX KO cell lines two individual clones from two independent sgRNAs were selected and expanded (KO1 and KO2) for further analyses. For the generation of ZNF274 KO cells, two independent sgRNAs were used; clones could only be obtained from one. One clone from each CAS9-only infection (V2) and non-specific sgRNA (random) were randomly chosen for controls. The four most likely off-target loci of each sgRNA were also sequenced to assure specificity KO with no off-target mutations observed in any of the clones (data not shown). For the generation of double KO clones, the ATRX KO2 clone was infected with lentiCRISPR V2 targeting ZNF274, selected, sorted and analyzed as above. In order to determine the allelespecific mutations, gDNA was extracted from each clone, amplified with specific primers to the target gRNA sequence (**Table S7**), cloned into the pCR4-TOPO vector, transformed into bacteria, and single bacterial colonies sequenced.

### Generation of DAXX KD cell lines by CRISP/Cas9

K562 cells were infected and selected as described above, with the exception that single cells were not sorted. Pools of KD cells were maintained in constant selection (puromycin 1ug/ml) and analyzed by immunoblotting to assess the efficiency of the KD. The double ATRX KO/DAXX KD was obtained by infecting and selecting ATRX KO2 cell line. Pools of cells infected with random sgRNA were used as control.

### Comet assay

The alkaline comet assay was performed as described with modifications. ^75^ K562 (1X10^5^) cells were washed with cold PBS, resuspended, and diluted in 100 uL of 0.5% low melting point agarose to be pipetted onto slides covered with 1.5% agarose. Cells were lysed (2.5M NaCl, 100 mM EDTA, 10mM Tris, pH 10, 1% Triton and 10% DMSO) for 24h at 4°C, incubated in electrophoresis buffer (300 mM NaOH, pH 13,1 mM EDTA) for 30 min and subjected to electrophoresis in the dark for 25 min at 25 V and 300 mA. Slides were neutralized 3x with Tris buffer (0.4M Tris, pH 7.5) for 5 min, dried with 100% ethanol and stained with ethidium bromide (20 ug/mL). Cells were imaged on a Nikon Eclipse^®^ microscope and ≥100 random cells were analyzed with CellProfiler software. Cell images were segmented using pixel intensity of 0.5 as threshold to generate masks matching the nucleoid. The comet tail was calculated by subtracting the nucleoid-integrated intensity from the comet-integrated intensity. For each sample, a positive control with cells treated with hydrogen peroxide (H_2_O_2_) (100uM for 30 min at 25°C) was analyzed concurrently. Experimental analysis was performed in a blinded fashion.

### Cell cycle analysis

K562 (3×10^6^) cells were pulsed with 10 uM of BrdU for 45 min, washed with PBS, and fixed in 70% cold ethanol for 1 hour on ice. The cells were then washed 2× with cold PBS and incubated in 2M hydrochloric acid for 30 min at room temperature. After incubation, cells were washed 2× with cold PBS and once with cold PBS-T solution (PBS, 0.2% Tween, 1% BSA), then stained with anti-BrdU FITC PBS-T (eBioscience, cat #11-5071-41) for 30 min in the dark. Cells were washed once with cold PBS-T, once more with cold PBS and then stained for 30 min on ice with a PI/RNAse solution (PBS, 20ug/ml PI, 10ug/ml RNAse). FACS analysis (10,000 cells per sample) was performed using BD FACS Canto II and FlowJo 10.

### H3K9me3 enrichment analysis

The significantly reduced H3K9me3 domains in the K562 ATRX KO2 H3K9me3 ChIP-seq were calculated with the SICER-df program. ^67^ Only regions overlapping significant H3K9me3 peaks in the V2 control were taken into account. H3K9me3 peaks in the H3K9me3 that were not significantly reduced compared to V2 were marked as unchanged. Bed files with the reduced and unchanged regions coordinates are provided in **Additional File 5**.

### Statistical analyses

The hypergeometric test and permutation tests implemented in the R coin package (v 1.0-24) were the primary statistical tests utilized (See **Table S2** for details). See **Table S3** for the results of all the performed tests.

### ChIP-qPCR and RT-qPCR

ChIP-qPCR and RT-qPCR were performed as described. ^76^ See **Table S7** for the list of primers used in our ChIP-qPCR and RT-qPCR experiments.

### Data Access

The datasets supporting the results of this article are available in the Gene Expression Omnibus (GEO) repository with the accession number GSE70920.

## AUTHOR’S CONTRIBUTIONS

DV-G and EB conceived this study with guidance and support from FR-T. DV-G and ZAQ performed ChIP-seq with assistance and support from DH and MAD. DV-G performed data analysis with help from DSM and ZAQ. DV-G derived the all KO cell lines and performed molecular characterization and cell cycle analysis with help from DSM. FGG performed comet assays and AHC performed co-IPs. DV-G and EB wrote the manuscript with assistance from ZAQ. All authors read and approved the final manuscript.

## ACKNOWLEDGMENTS

The authors thank the Genomics Core Facility, the Flow Cytometry Center, the Department of Scientific Computing and the Pluripotent Stem Cell Core Facility at Mount Sinai for technical assistance. We thank Matthew O'Connell and Kajan Ratnakumar for helpful discussions and guidance. Funding was provided by the DGAPA-PAPIIT, UNAM (IN209403, IN203811 and IN201114), CONACyT (42653-Q, 128464 and 220503) and Fronteras de la Ciencia-2015 CONACyT (Grant 290) to FR-T, Graduate fellowship from CONACyT (239663, CVU 257385) to DV-G, NCI T32-CA078207 to ZAQ, NIH EY014867, EY018599 and CA168875, Cancer Center Support from the NCI (CA21765), support from the American Lebanese Syrian Associated Charities (ALSAC) and a grant from Alex’s Lemonade Stand Foundation for Childhood Cancer to MAD, and NCI/NIH R01CA154683 to EB. This article is part of the doctoral thesis of DV-G from the Doctorado en Ciencias Biomédicas, UNAM.

## FIGURE LEGENDS

**Supplementary Figure S1.**
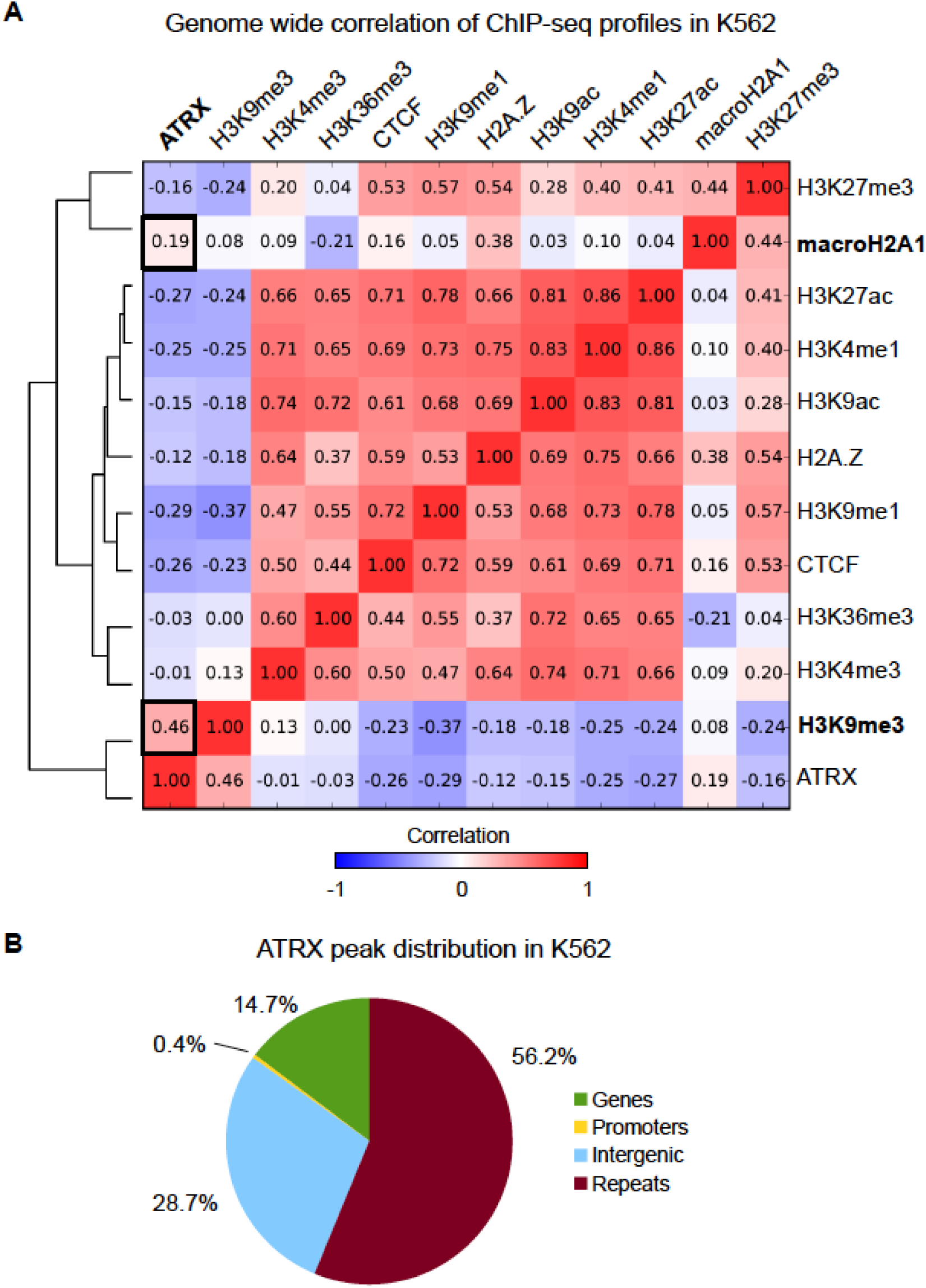
ATRX correlates with H3K9me3 genome-wide <u>in K562 cells</u>. (A) Correlation plot of ATRX ChIP-seq enrichment with re-analyzed ENCODE datasets for histone modifications, TFs and histone variants in K562 cells. (B) Distribution of ATRX significant peaks in genes (TSS-TES), promoters (-3kb to TSS), intergenic and repetitive (regions masked by repeat masker) sequences <u>in K562</u>.

**Supplementary Figure S2.**
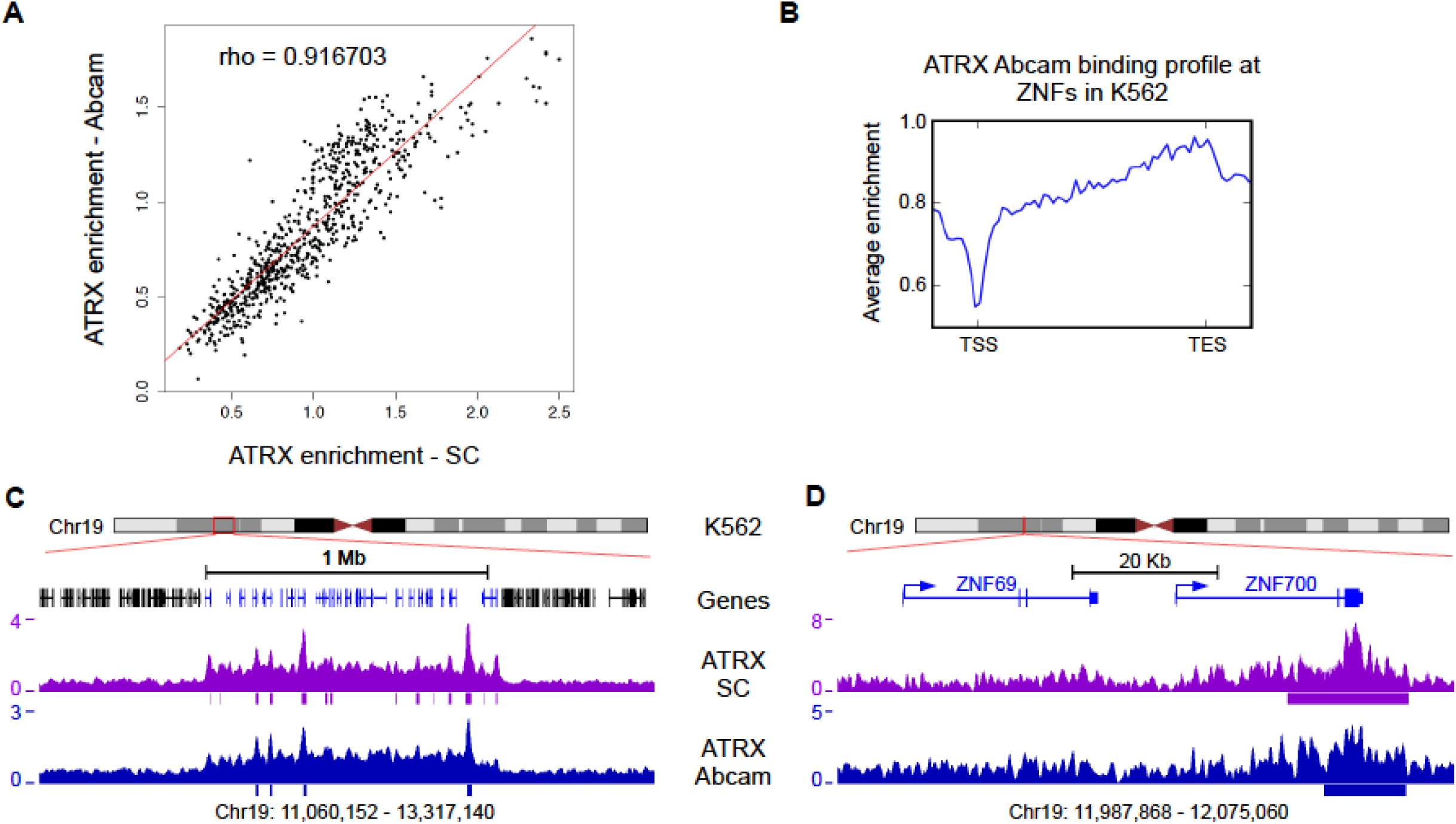
Independent antibodies for ATRX ChIP-seq analysis show similar enrichment patterns <u>in K562 cells</u>. (A) Correlation of enrichment over ZNF genes for two independent antibodies <u>in K562</u>. Each dot represents one ZNF gene. X axis = average enrichment over input for the ATRX Santa Cruz (SC) antibody; Y axis = average enrichment over input for the ATRX Abcam antibody. Antibodies show a high enrichment correlation. (B) Average enrichment ChIP-seq profile of ATRX Abcam over all ZNF gene bodies +/- 1kb shows enrichment at the 3’ region <u>in K562</u>. The pattern is very similar to the one observed for ATRX SC (see Fig. 1G, left). (C) UCSC Genome browser screenshot of a typical ZNF cluster in chromosome 19 (blue genes). The enrichment over input signals for both ATRX antibodies are shown. Significant peaks are shown as bars below enrichment tracks. (D) Zoomed in snapshot of two genes contained in the ZNF cluster shown in (C).

**Supplementary Figure S3.**
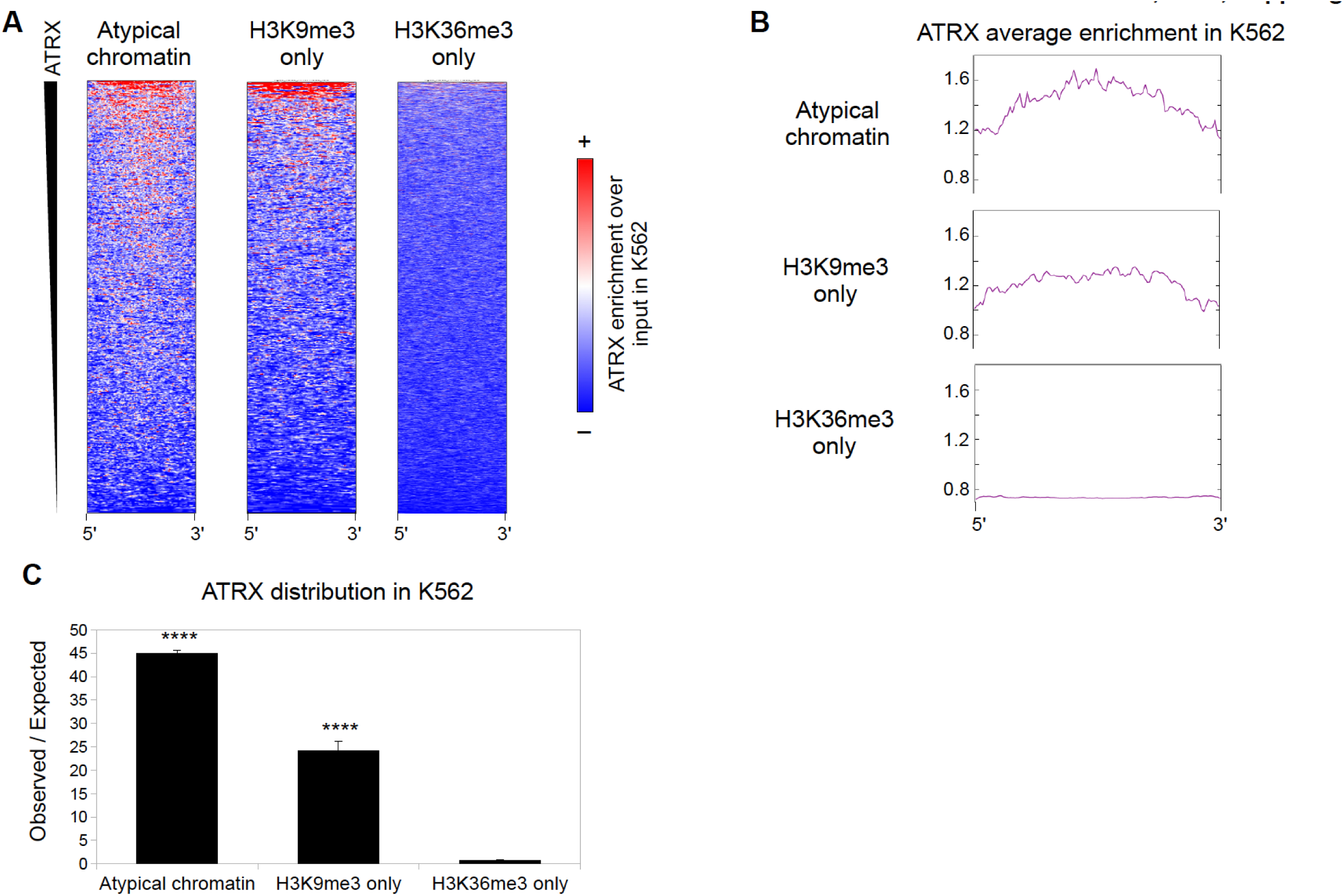
ATRX is enriched at atypical chromatin regions genome-wide <u>in K562 cells</u>. (A) Metagene plots of ATRX enrichment over input in atypical chromatin (left), H3K9me3-only (middle) and H3K36me3-only (right) peaks. The regions are sorted by ATRX enrichment from high to low from top to down, respectively. (B) Average enrichment ChIP-seq profile from the plots shown in (A). (C) Observed over expected enrichment of ATRX peaks in atypical, H3K9me3-only and H3K36me3-only regions <u>in K562</u>.

**Supplementary Figure S4.**
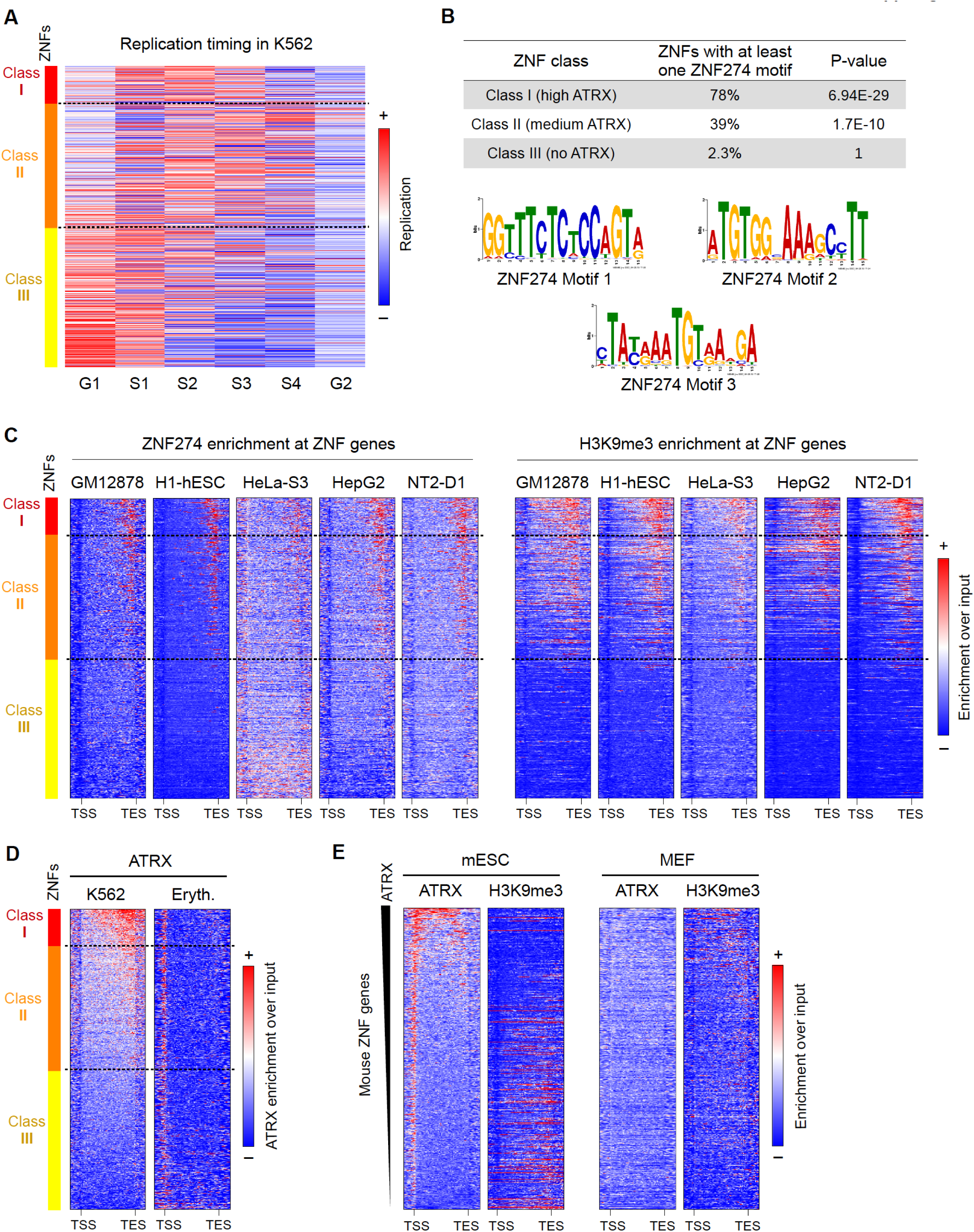
ATRX-bound ZNFs show particular genetic and epigenetic features. (A) <u>K562</u> Repli-seq average enrichments over ZNFs at distinct points of the cell cycle. S1 and S2 are early S phase, whereas S3 and S4 represent the late stages of S phase. Genes are sorted by Class (ATRX enrichment from high to low, top to bottom, respectively). (B) Enrichment of ZNF274 binding motifs in different ZNF Classes (top). Logos of the three identified ZNF274 binding motifs (bottom). (C) Metagene plots over ZNFs of ZNF274 and H3K9me3 in different cell lines show that the pattern observed in K562 is maintained in most cell lines. The genes are sorted as in (A). (D) ATRX ChIP-seq metagene profiles of ATRX enrichment over ZNFs in K562 cells and human erythroblasts. (E) ATRX and H3K9me3 ChIP-seq metagene profiles over ZNFs in two mouse cell types. Mouse ZNF genes were sorted from top to bottom by their ATRX enrichment in Embryonic Stem Cells (mESC) from high to low. See Table S6 for ChIP-seq data sources.

**Supplementary Figure S5.**
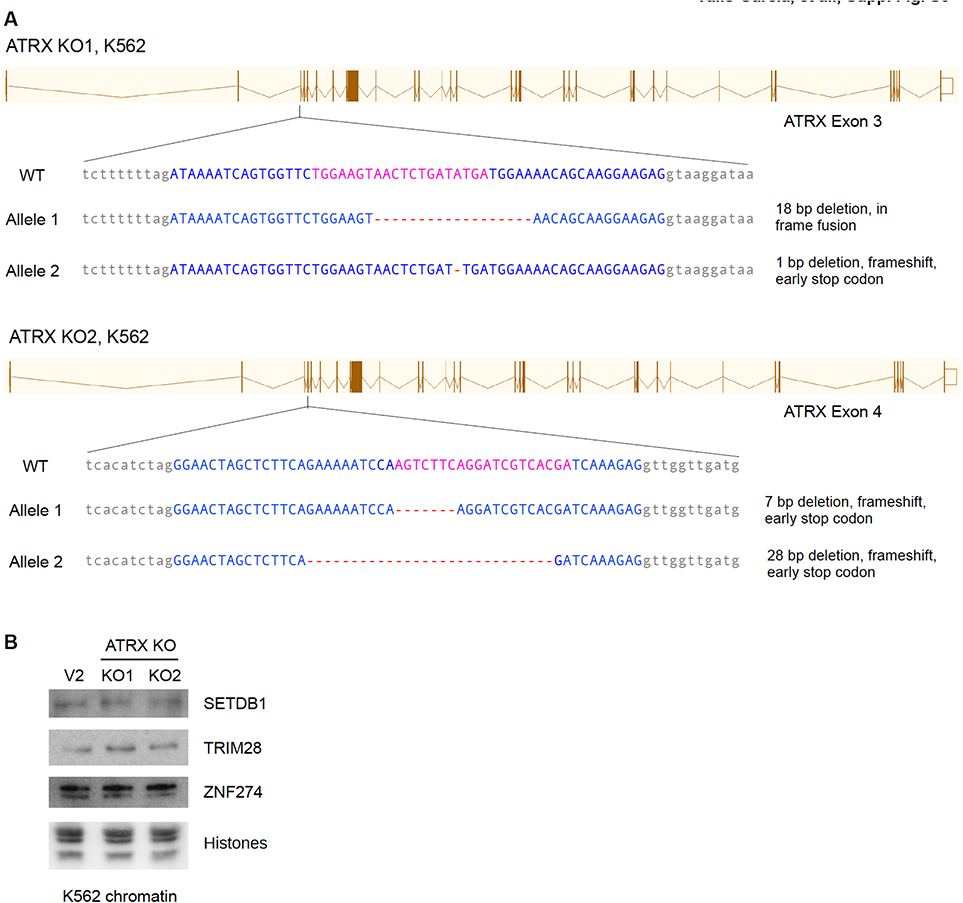
Characterization of ATRX KO K562 cell lines. (A) Mutations observed in the ATRX KO <u>K562</u> cell lines. The brown diagrams show the ATRX gene with exons as boxes and introns as lines. The black lines show the zoom in to the exons that are mutated in the KO1 cell line (top) and the KO2 (bottom) cell line. The WT sequence is shown as comparison to the KO sequences. The pink bases in the WT depict the region targeted by the sgRNA, the red dashes show the bases that were deleted in the mutant clones of ATRX. Each allele indicates the type of mutation and the consequence at the protein level. Blue upper case bases are exonic; grey lower case bases are intronic. (B) Chromatin immunoblots in control V2 and ATRX KO <u>K562</u> cell lines of proteins that interact with ATRX. Amido black staining of histones is shown for loading.

**Supplementary Figure S6.**
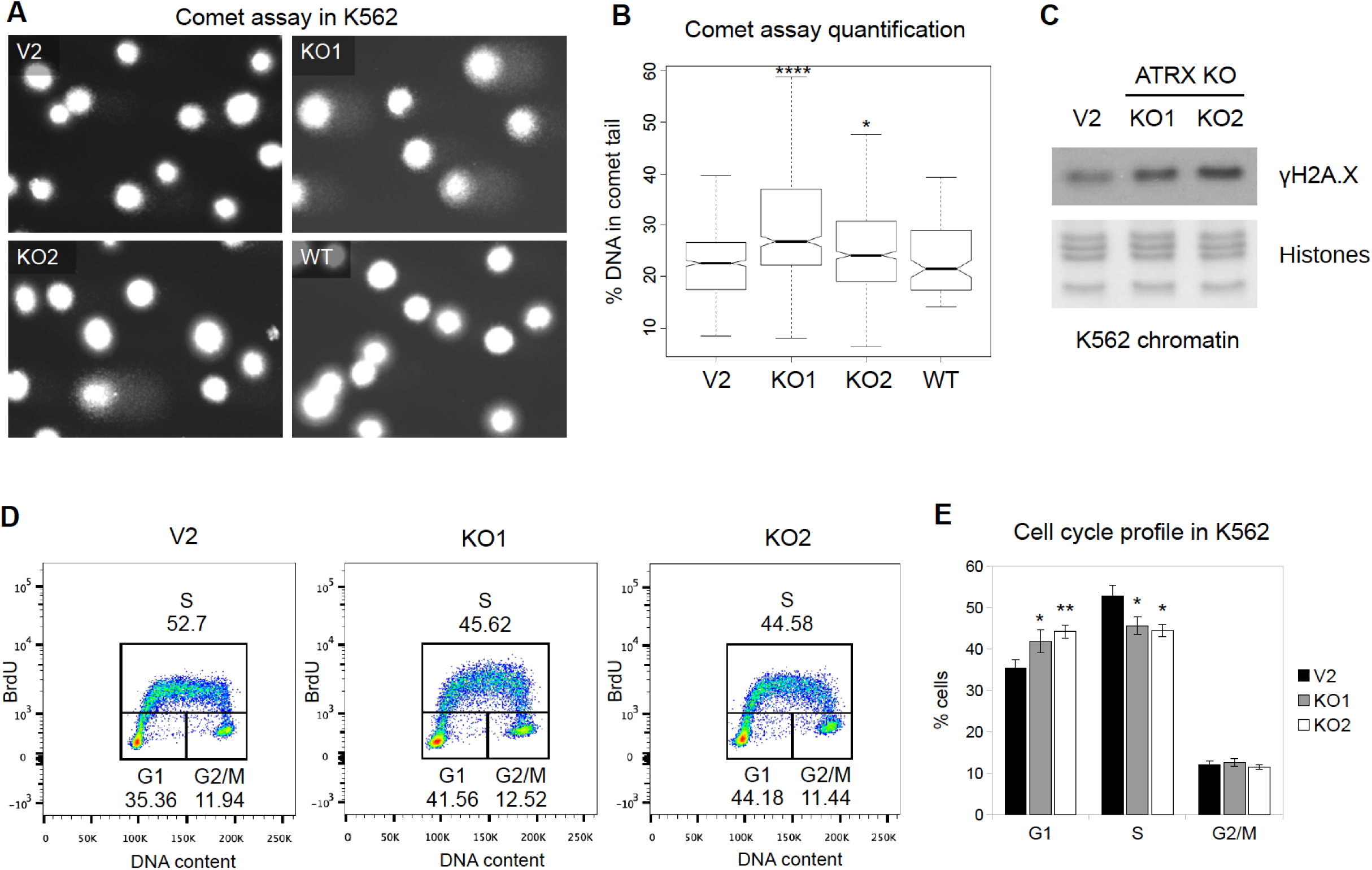
ATRX KO K562 cells have higher percentage of DNA damage and defects in cell cycle. (A) Representative slides of the comet assay performed in WT, control V2, ATRX KO1 and ATRX KO2 <u>K562</u> cells. (B) Quantification of 4 independent comet assay replicates (>100 cells analyzed per replicate). Asterisks show significant differences compared to the V2 control (* pvalue < 0.05; **** pvalue < 1×10-4). (C) Chromatin immunoblot of γH2A.X in ATRX KO <u>K562</u> cells. Histones are shown as a loading control. (D) Representative <u>K562</u> cell cycle profiles of control V2 and ATRX KO1 and KO2 cells assessed by BrdU/PI staining profiles. The numbers show the average of at least 5 independent experiments. (E) Quantification of at least 5 independent BrdU/PI profile replicates. The bars show the average % of cells in each phase, error bars depict SEM. Asterisks show significant changes compared to the control V2 (* pvalue < 0.05; ** pvalue < 0.01).

**Supplementary Figure S7.**
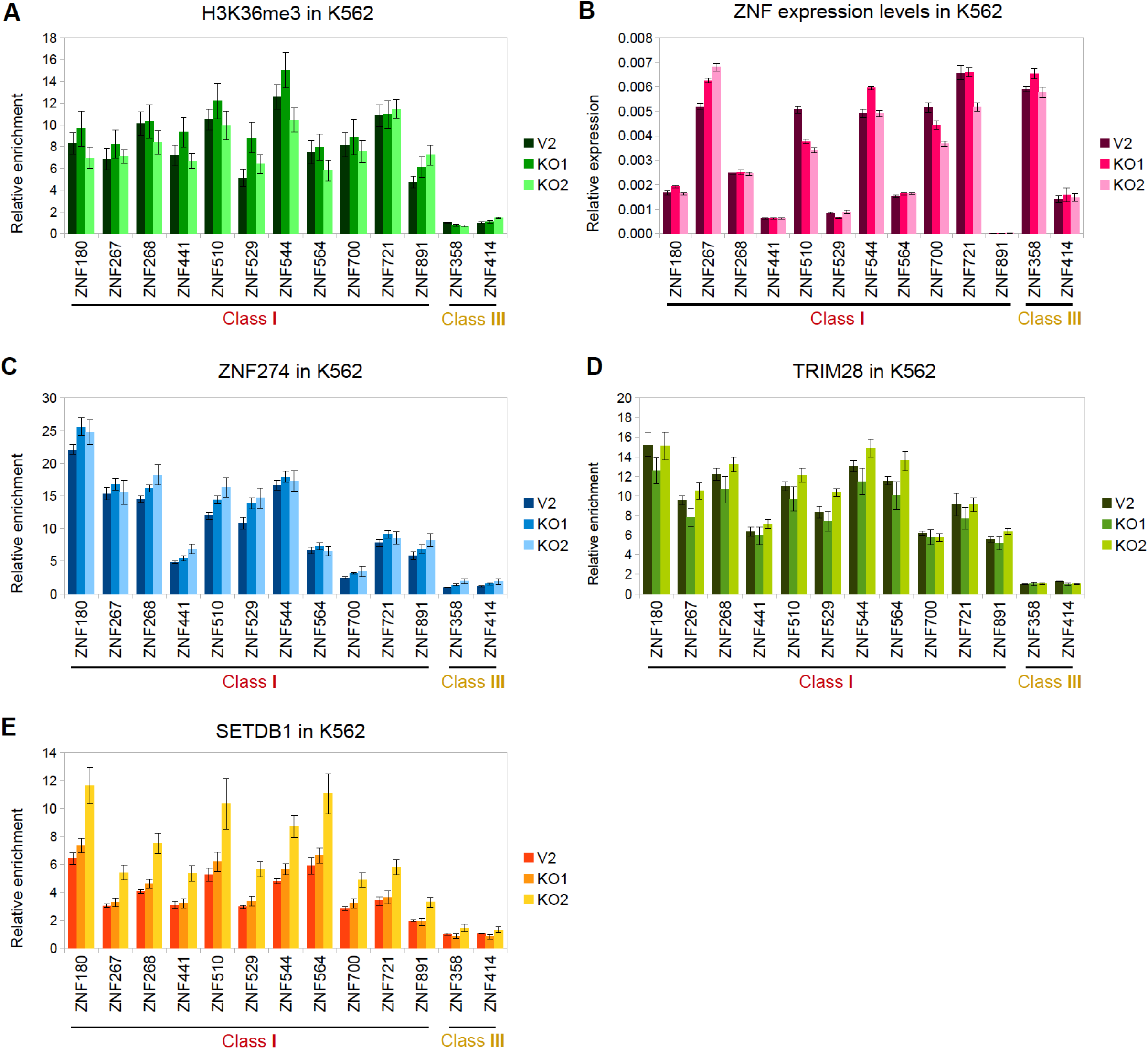
Effect of ATRX KO at ZNFs <u>in K562 cells</u>. (A) Native ChIP-qPCR of H3K36me3 over the 3’ exon region of ZNF genes in control V2 and ATRX KO <u>K562</u> cell lines. (B) RT-qPCR of the panel of ZNFs from (A) in V2, KO1 and KO2 cells. (C-E) ChIP-qPCR of ZNF274 (C), TRIM28 (D) and SETDB1 (E) in V2, KO1 and KO2 cell lines using the same ZNF panel as in (A). All graphs show the average of at least three independent biological replicates. Error bars represent SEM. For details about statistical comparisons see **Table S3**.

**Supplementary Figure S8.**
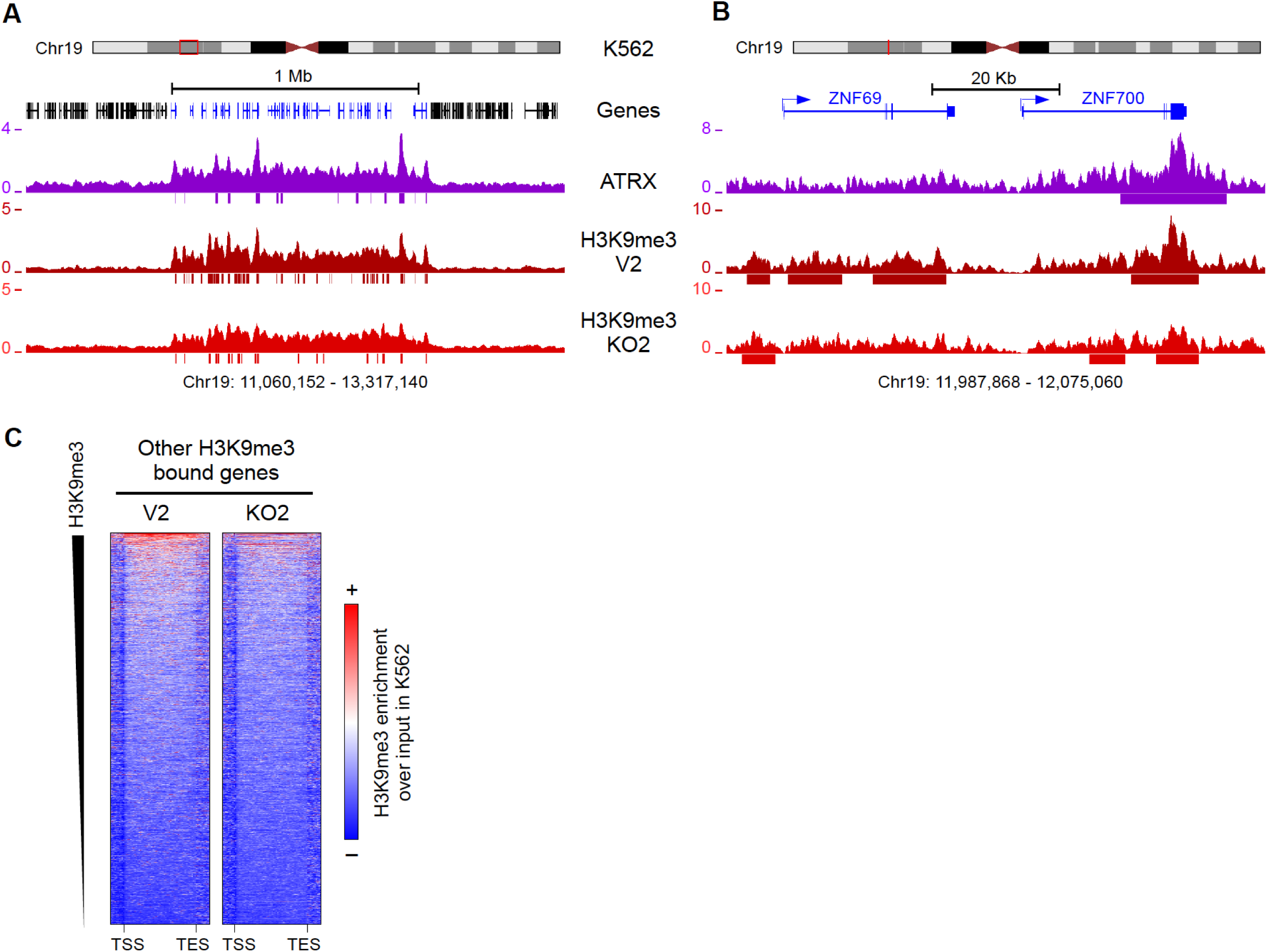
H3K9me3 loss in ATRX KO <u>K562</u> cell line. (A) UCSC genome browser screenshot of a typical ZNF cluster in chromosome 19 (blue genes). The enrichment over input signals for ATRX and H3K9me3 in control V2 and <u>ATRX</u> KO2 cells is shown. Significant peaks are shown as bars below the enrichment tracks. (B) Zoomed in screenshot of two genes contained in the ZNF cluster shown in (A). (C) Metagene plot of H3K9me3 ChIP-seq enrichment in H3K9me3-bound genes in control V2 and ATRX KO2 cells. The genes are ordered by their H3K9me3 enrichment from high to low from top to bottom, respectively.

**Supplementary Figure S9.**
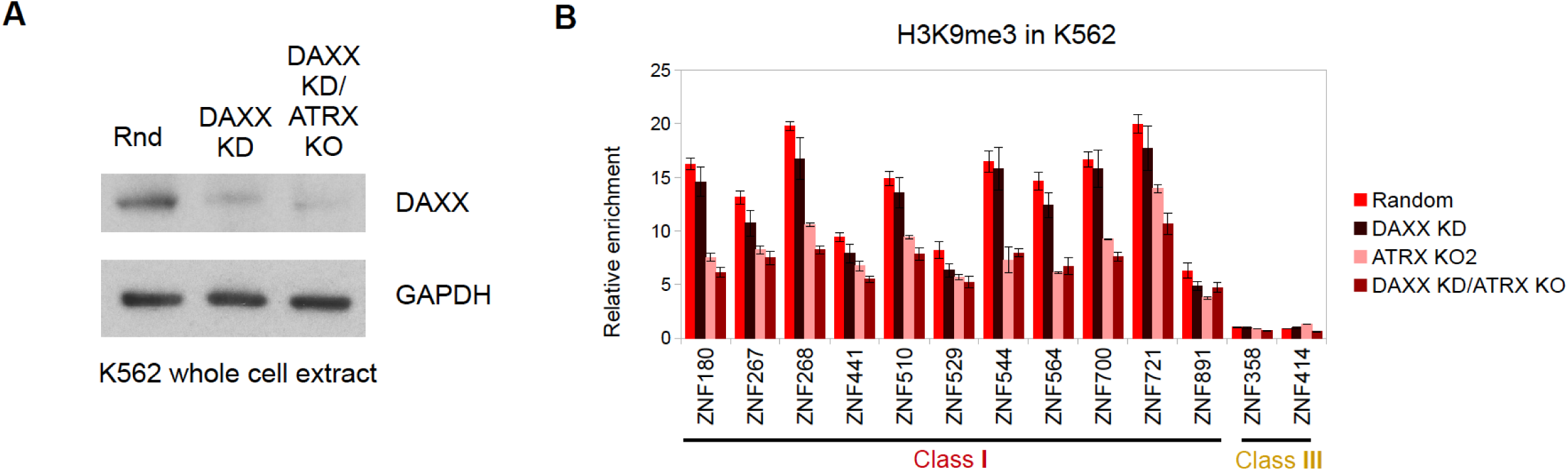
DAXX KD does not affect H3K9me3 levels at ZNF genes <u>in K562 cells</u>. (A) Immunoblot of <u>K562</u> whole cell extracts in control (Rnd) and DAXX single KD and double DAXX KD/ATRX KO cells. GAPDH is used as a loading control. (B) H3K9me3 ChIP-qPCR over ZNF genes in control and DAXX KD and ATRX KO single and double mutant <u>K562</u> cells. Bars show the average of at least two biological replicates; the error bars depict SEM.

**Supplementary Figure S10.**
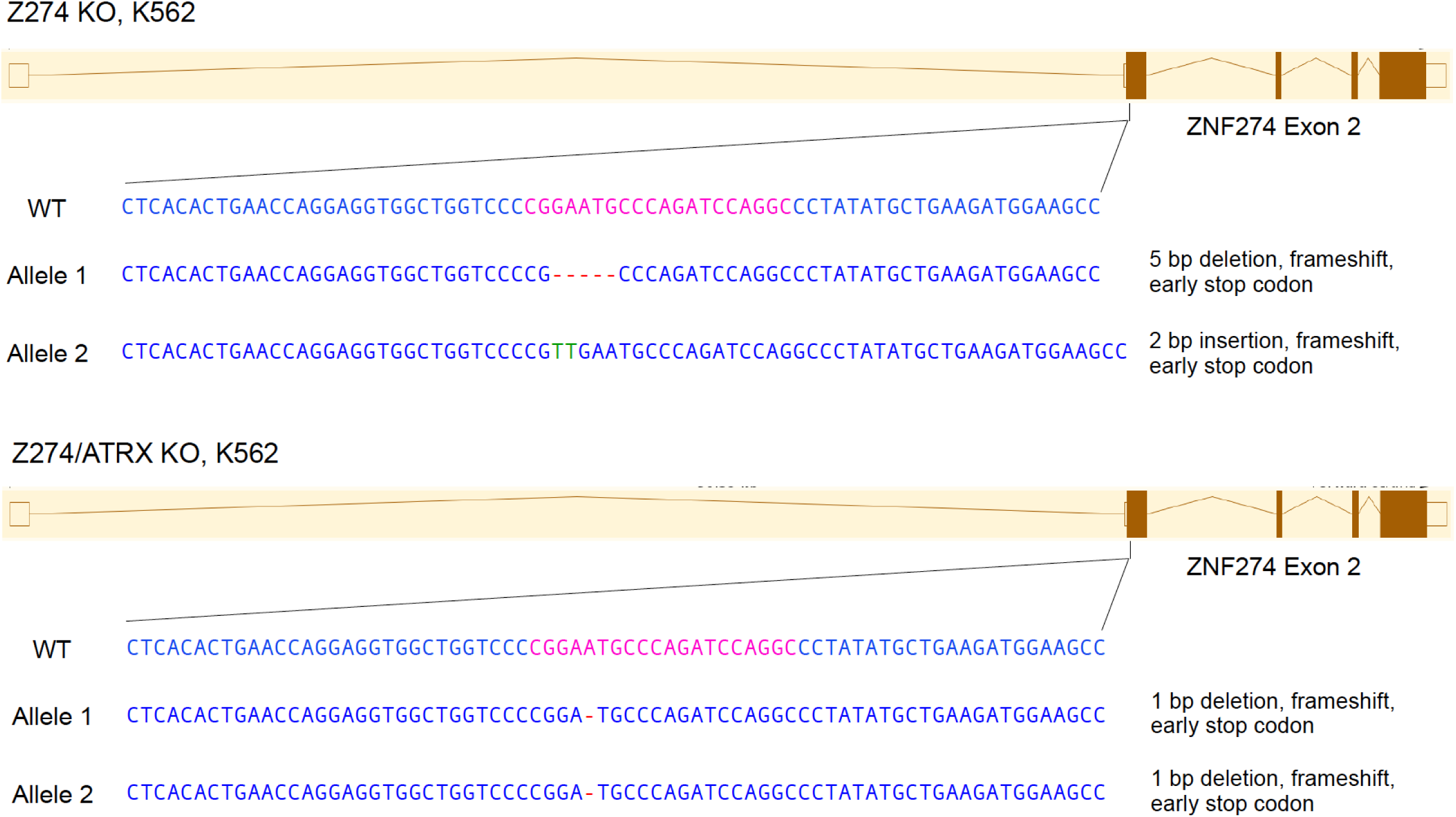
Characterization of ZNF274 KO <u>K562</u> cell lines. Mutations observed in the ZNF274 KO <u>K562</u> cell lines. The brown diagrams show the ZNF274 gene with exons as boxes and introns as lines. The black lines show the zoom in to the exon that is mutated in the single ZNF274 (Z274) KO cell line (top) and the double ZNF274 * ATRX KO (bottom, Z274/ATRX KO) cell line. The WT sequence is shown as comparison to the KO sequences. The pink bases in the WT depict the region targeted by the sgRNA, the red dashes show the bases that were deleted in the mutant clones of ATRX. Green bases indicate insertions. Each allele indicates the type of mutation and the consequence at the protein level.

## REFERENCES

1. Sinha M, Peterson CL. Chromatin dynamics during repair of chromosomal DNA double-strand breaks. Epigenomics 2009; 1:371–85.

2. Hargreaves DC, Crabtree GR. ATP-dependent chromatin remodeling: genetics, genomics and mechanisms. Cell Res 2011; 21:396–420.

3. Euskirchen G, Auerbach RK, Snyder M. SWI/SNF chromatin-remodeling factors: Multiscale analyses and diverse functions. J Biol Chem 2012; 287:30897–905.

4. Gospodinov A, Herceg Z. Chromatin structure in double strand break repair. DNA Repair (Amst) 2013; 12:800–10.

5. Längst G, Manelyte L. Chromatin Remodelers: From Function to Dysfunction. Genes (Basel) 2015; 6:299–324.

6. Clynes D, Gibbons RJ. ATRX and the replication of structured DNA. Curr Opin Genet Dev 2013; 23:289–94.

7. Voon HPJ, Wong LH. New players in heterochromatin silencing: histone variant H3.3 and the ATRX/DAXX chaperone. Nucleic Acids Res 2016;:gkw012.

8. Ratnakumar K, Bernstein E. ATRX: the case of a peculiar chromatin remodeler. Epigenetics 2013; 8:3–9.

9. Watson LA, Goldberg H, Bérubé NG. Emerging roles of ATRX in cancer. Epigenomics 2015; 7:1365–78.

10. Park DJ, Pask AJ, Huynh K, Renfree MB, Harley VR, Marshall Graves J a. Comparative analysis of ATRX, a chromatin remodeling protein. Gene 2004; 339:39–48.

11. Eustermann S, Yang J-C, Law MJ, Amos R, Chapman LM, Jelinska C, Garrick D, Clynes D, Gibbons RJ, Rhodes D, et al. Combinatorial readout of histone H3 modifications specifies localization of ATRX to heterochromatin. Nat Struct Mol Biol 2011; 18:777–82.

12. Iwase S, Xiang B, Ghosh S, Ren T, Lewis PW, Cochrane JC, Allis CD, Picketts DJ, Patel DJ, Li H, et al. ATRX ADD domain links an atypical histone methylation recognition mechanism to human mental-retardation syndrome. Nat Struct Mol Biol 2011; 18:769–76.

13. Xue Y, Gibbons R, Yan Z, Yang D, McDowell TL, Sechi S, Qin J, Zhou S, Higgs D, Wang W. The ATRX syndrome protein forms a chromatin-remodeling complex with Daxx and localizes in promyelocytic leukemia nuclear bodies. Proc Natl Acad Sci U S A 2003; 100:10635–40.

14. McDowell TL, Gibbons RJ, Sutherland H, O’Rourke DM, Bickmore W a, Pombo a, Turley H, Gatter K, Picketts DJ, Buckle VJ, et al. Localization of a putative transcriptional regulator (ATRX) at pericentromeric heterochromatin and the short arms of acrocentric chromosomes. Proc Natl Acad Sci U S A 1999; 96:13983–8.

15. Law MJ, Lower KM, Voon HPJJ, Hughes JR, Garrick D, Viprakasit V, Mitson M, De Gobbi M, Marra M, Morris A, et al. ATR-X Syndrome Protein Targets Tandem Repeats and Influences Allele-Specific Expression in a Size-Dependent Manner. Cell 2010; 143:367–78.

16. Drané P, Ouararhni K, Depaux A, Shuaib M, Hamiche A. The death-associated protein DAXX is a novel histone chaperone involved in the replication-independent deposition of H3.3. Genes Dev 2010; 24:1253–65.

17. Goldberg AD, Banaszynski L a, Noh K-M, Lewis PW, Elsaesser SJ, Stadler S, Dewell S, Law M, Guo X, Li X, et al. Distinct factors control histone variant H3.3 localization at specific genomic regions. Cell 2010; 140:678–91.

18. Wong LH, McGhie JD, Sim M, Anderson M a, Ahn S, Hannan RD, George AJ, Morgan K a, Mann JR, Choo KHA. ATRX interacts with H3.3 in maintaining telomere structural integrity in pluripotent embryonic stem cells. Genome Res 2010; 20:351–60.

19. Lewis PW, Elsaesser SJ, Noh K-M, Stadler SC, Allis CD. Daxx is an H3.3-specific histone chaperone and cooperates with ATRX in replication-independent chromatin assembly at telomeres. Proc Natl Acad Sci U S A 2010; 107:14075–80.

20. Sadic D, Schmidt K, Groh S, Kondofersky I, Ellwart J, Fuchs C, Theis FJ, Schotta G. Atrx promotes heterochromatin formation at retrotransposons. EMBO Rep 2015;:1–15.

21. Elsässer SJ, Noh K-M, Diaz N, Allis CD, Banaszynski L a. Histone H3.3 is required for endogenous retroviral element silencing in embryonic stem cells. Nature 2015;

22. He Q, Kim H, Huang R, Lu W, Tang M, Shi F, Yang D, Zhang X, Huang J, Liu D, et al. The Daxx/Atrx Complex Protects Tandem Repetitive Elements during DNA Hypomethylation by Promoting H3K9 Trimethylation. Cell Stem Cell 2015; 17:273–86.

23. Bérubé NG, Smeenk C a, Picketts DJ. Cell cycle-dependent phosphorylation of the ATRX protein correlates with changes in nuclear matrix and chromatin association. Hum Mol Genet 2000; 9:539–47.

24. Lechner MS, Schultz DC, Negorev D, Maul GG, Rauscher FJ. The mammalian heterochromatin protein 1 binds diverse nuclear proteins through a common motif that targets the chromoshadow domain. Biochem Biophys Res Commun 2005; 331:929–37.

25. Voon HPJ, Hughes JR, Rode C, De La Rosa-Velázquez IA, Jenuwein T, Feil R, Higgs DR, Gibbons RJ. ATRX Plays a Key Role in Maintaining Silencing at Interstitial Heterochromatic Loci and Imprinted Genes. Cell Rep 2015;:405–18.

26. Ray-Gallet D, Woolfe A, Vassias I, Pellentz C, Lacoste N, Puri A, Schultz DC, Pchelintsev N a., Adams PD, Jansen LET, et al. Dynamics of Histone H3 Deposition In vivo Reveal a Nucleosome Gap-Filling Mechanism for H3.3 to Maintain Chromatin Integrity. Mol Cell 2011; 44:928–41.

27. Schneiderman JI, Orsi G a, Hughes KT, Loppin B, Ahmad K. Nucleosome-depleted chromatin gaps recruit assembly factors for the H3.3 histone variant. Proc Natl Acad Sci U S A 2012; 109:19721–6.

28. Filipescu D, Szenker E, Almouzni G. Developmental roles of histone H3 variants and their chaperones. Trends Genet 2013; 29:630–40.

29. Ratnakumar K, Duarte LF, LeRoy G, Hasson D, Smeets D, Vardabasso C, Bönisch C, Zeng T, Xiang B, Zhang DY, et al. ATRX-mediated chromatin association of histone variant macroH2A1 regulates α-globin expression. Genes Dev 2012; 26:433–8.

30. Clynes D, Higgs DR, Gibbons RJ. The chromatin remodeller ATRX: a repeat offender in human disease. Trends Biochem Sci 2013; 38:461–6.

31. Levy M a., Kernohan KD, Jiang Y, Bérubé NG. ATRX promotes gene expression by facilitating transcriptional elongation through guanine-rich coding regions. Hum Mol Genet 2015; 24:1824–35.

32. Heaphy CM, de Wilde RF, Jiao Y, Klein AP, Edil BH, Shi C, Bettegowda C, Rodriguez FJ, Eberhart CG, Hebbar S, et al. Altered telomeres in tumors with ATRX and DAXX mutations. Science 2011; 333:425.

33. Schwartzentruber J, Korshunov A, Liu X-Y, Jones DTW, Pfaff E, Jacob K, Sturm D, Fontebasso AM, Quang D-AK, Tönjes M, et al. Driver mutations in histone H3.3 and chromatin remodelling genes in paediatric glioblastoma. Nature 2012; 484:130–130.

34. Lovejoy C a, Li W, Reisenweber S, Thongthip S, Bruno J, de Lange T, De S, Petrini JHJ, Sung P a, Jasin M, et al. Loss of ATRX, genome instability, and an altered DNA damage response are hallmarks of the alternative lengthening of telomeres pathway. PLoS Genet 2012; 8:e1002772.

35. Bower K, Napier CE, Cole SL, Dagg R a, Lau LMS, Duncan EL, Moy EL, Reddel RR. Loss of wild-type ATRX expression in somatic cell hybrids segregates with activation of Alternative Lengthening of Telomeres. PLoS One 2012; 7:e50062.

36. Clynes D, Jelinska C, Xella B, Ayyub H, Scott C, Mitson M, Taylor S, Higgs DR, Gibbons RJ. Suppression of the alternative lengthening of telomere pathway by the chromatin remodelling factor ATRX. Nat Commun 2015; 6:7538.

37. Huh MS, Price O’Dea T, Ouazia D, McKay BC, Parise G, Parks RJ, Rudnicki MA, Picketts DJ. Compromised genomic integrity impedes muscle growth after Atrx inactivation. J Clin Invest 2012; 122:4412–23.

38. Watson LA, Solomon L a., Li JR, Jiang Y, Edwards M, Shin-Ya K, Beier F, Bérubé NG. Atrx deficiency induces telomere dysfunction, endocrine defects, and reduced life span. J Clin Invest 2013; 123:2049–63.

39. Leung JW-CC, Ghosal G, Wang W, Shen X, Wang J, Li L, Chen J. Alpha thalassemia/mental retardation syndrome X-linked gene product ATRX is required for proper replication restart and cellular resistance to replication stress. J Biol Chem 2013; 288:6342–50.

40. Clynes D, Jelinska C, Xella B, Ayyub H, Taylor S, Mitson M, Bachrati CZ, Higgs DR, Gibbons RJ. ATRX dysfunction induces replication defects in primary mouse cells. PLoS One 2014; 9:e92915.

41. Emerson RO, Thomas JH. Adaptive evolution in zinc finger transcription factors. PLoS Genet 2009; 5.

42. Dhwani Tadepally H, Aubry M. Evolution of C2H2 Zinc-finger Gene Families in Mammals. In: Encyclopedia of Life Sciences. Chichester, UK: John Wiley & Sons, Ltd; 2010. page 1–9.

43. Nowick K, Fields C, Gernat T, Caetano-Anolles D, Kholina N, Stubbs L. Gain, loss and divergence in primate zinc-finger genes: a rich resource for evolution of gene regulatory differences between species. PLoS One 2011; 6:e21553.

44. Blahnik KR, Dou L, Echipare L, Iyengar S, O’Geen H, Sanchez E, Zhao Y, Marra M a, Hirst M, Costello JF, et al. Characterization of the contradictory chromatin signatures at the 3’ exons of zinc finger genes. PLoS One 2011; 6:e17121.

45. Dunham I, Kundaje A, Aldred SF, Collins PJ, Davis C a, Doyle F, Epstein CB, Frietze S, Harrow J, Kaul R, et al. An integrated encyclopedia of DNA elements in the human genome. Nature 2012; 489:57–74.

46. Sarma K, Cifuentes-Rojas C, Ergun A, Del Rosario A, Jeon Y, White F, Sadreyev R, Lee JT. ATRX directs binding of PRC2 to Xist RNA and Polycomb targets. Cell 2014; 159:869–83.

47. Ramamoorthy M, Smith S. Loss of ATRX Suppresses Resolution of Telomere Cohesion to Control Recombination in ALT Cancer Cells. Cancer Cell 2015; 28:357–69.

48. Ernst J, Kheradpour P, Mikkelsen TS, Shoresh N, Ward LD, Epstein CB, Zhang X, Wang L, Issner R, Coyne M, et al. Mapping and analysis of chromatin state dynamics in nine human cell types. Nature 2011; 473:43–9.

49. Vogel MJ, Guelen L, De Wit E, Peric-Hupkes D, Lodén M, Talhout W, Feenstra M, Abbas B, Classen AK, Van Steensel B. Human heterochromatin proteins form large domains containing KRAB-ZNF genes. Genome Res 2006; 16:1493–504.

50. Julienne H, Zoufir A, Audit B, Arneodo A. Human genome replication proceeds through four chromatin states. PLoS Comput Biol 2013; 9:e1003233.

51. Hoffman MM, Ernst J, Wilder SP, Kundaje A, Harris RS, Libbrecht M, Giardine B, Ellenbogen PM, Bilmes J a, Birney E, et al. Integrative annotation of chromatin elements from ENCODE data. Nucleic Acids Res 2013; 41:827–41.

52. Iyengar S, Farnham PJ. KAP1 protein: An enigmatic master regulator of the genome. J Biol Chem 2011; 286:26267–76.

53. Frietze S, O’Geen H, Blahnik KR, Jin VX, Farnham PJ. ZNF274 recruits the histone methyltransferase SETDB1 to the 3’ ends of ZNF genes. PLoS One 2010; 5:e15082.

54. Bérubé NG. ATRX in chromatin assembly and genome architecture during development and disease. Biochem Cell Biol 2011; 89:435–44.

55. Najafabadi HS, Mnaimneh S, Schmitges FW, Garton M, Lam KN, Yang A, Albu M, Weirauch MT, Radovani E, Kim PM, et al. C2H2 zinc finger proteins greatly expand the human regulatory lexicon. Nat Biotechnol 2015; 33:555–62.

56. Iyengar S, Ivanov A V, Jin VX, Rauscher FJ, Farnham PJ. Functional analysis of KAP1 genomic recruitment. Mol Cell Biol 2011; 31:1833–47.

57. Aymard F, Bugler B, Schmidt CK, Guillou E, Caron P, Briois S, Iacovoni JS, Daburon V, Miller KM, Jackson SP, et al. Transcriptionally active chromatin recruits homologous recombination at DNA double-strand breaks. Nat Struct Mol Biol 2014; 21:366–74.

58. Pfister SX, Ahrabi S, Zalmas L-P, Sarkar S, Aymard F, Bachrati CZ, Helleday T, Legube G, La Thangue NB, Porter ACG, et al. SETD2-dependent histone H3K36 trimethylation is required for homologous recombination repair and genome stability. Cell Rep 2014; 7:2006–18.

59. Murray JM, Stiff T, Jeggo P a. DNA double-strand break repair within heterochromatic regions. Biochem Soc Trans 2012; 40:173–8.

60. Kalousi A, Hoffbeck A-S, Selemenakis PN, Pinder J, Savage KI, Khanna KK, Brino L, Dellaire G, Gorgoulis VG, Soutoglou E. The Nuclear Oncogene SET Controls DNA Repair by KAP1 and HP1 Retention to Chromatin. Cell Rep 2015; 11:149–63.

61. Napier CE, Huschtscha LI, Harvey A, Bower K, Noble JR, Hendrickson E a, Reddel RR. ATRX represses alternative lengthening of telomeres. Oncotarget 2015; 6:16543–58.

62. Naftelberg S, Schor IE, Ast G, Kornblihtt AR. Regulation of Alternative Splicing Through Coupling with Transcription and Chromatin Structure. Annu Rev Biochem 2015; 84:165–98.

63. Hasson D, Panchenko T, Salimian KJ, Salman MU, Sekulic N, Alonso A, Warburton PE, Black BE. The octamer is the major form of CENP-A nucleosomes at human centromeres. Nat Struct Mol Biol 2013; 20:687–95.

64. Langmead B, Trapnell C, Pop M, Salzberg SL. Ultrafast and memory-efficient alignment of short DNA sequences to the human genome. Genome Biol 2009; 10:R25.

65. Zhang Y, Liu T, Meyer C a, Eeckhoute J, Johnson DS, Bernstein BE, Nusbaum C, Myers RM, Brown M, Li W, et al. Model-based analysis of ChIP-Seq (MACS). Genome Biol 2008; 9:R137.

66. Liang K, Keleş S. Normalization of ChIP-seq data with control. BMC Bioinformatics 2012; 13:199.

67. Zang C, Schones DE, Zeng C, Cui K, Zhao K, Peng W. A clustering approach for identification of enriched domains from histone modification ChIP-Seq data. Bioinformatics 2009; 25:1952–8.

68. Quinlan AR, Hall IM. BEDTools: A flexible suite of utilities for comparing genomic features. Bioinformatics 2010; 26:841–2.

69. Ramírez F, Dündar F, Diehl S, Grüning B a., Manke T. DeepTools: A flexible platform for exploring deep-sequencing data. Nucleic Acids Res 2014; 42:187–91.

70. Sandve GK, Gundersen S, Johansen M, Glad IK, Gunathasan K, Holden L, Holden M, Liestøl K, Nygård S, Nygaard V, et al. The Genomic HyperBrowser: an analysis web server for genome-scale data. Nucleic Acids Res 2013; 41:W133–41.

71. Huang DW, Sherman BT, Lempicki R a. Systematic and integrative analysis of large gene lists using DAVID bioinformatics resources. Nat Protoc 2009; 4:44–57.

72. Huppert JL, Balasubramanian S. Prevalence of quadruplexes in the human genome. Nucleic Acids Res 2005; 33:2908–16.

73. Machanick P, Bailey TL. MEME-ChIP: Motif analysis of large DNA datasets. Bioinformatics 2011; 27:1696–7.

74. Ran F, Hsu P, Wright J, Agarwala V. Genome engineering using the CRISPRCas9 system. Nat Protoc 2013; 8:2281–308.

75. Singh NP, McCoy MT, Tice RR, Schneider EL. A simple technique for quantitation of low levels of DNA damage in individual cells. Exp Cell Res 1988; 175:184–91.

76. Vardabasso C, Gaspar-Maia A, Hasson D, Pünzeler S, Valle-Garcia D, Straub T, Keilhauer EC, Strub T, Dong J, Panda T, et al. Histone Variant H2A.Z.2 Mediates Proliferation and Drug Sensitivity of Malignant Melanoma. Mol Cell 2015;:75–88.

